# Redox Dyshomeostasis Links Renal and Neuronal Dysfunction in *Drosophila* Models of Gaucher and Parkinson’s Disease

**DOI:** 10.1101/2025.08.06.668868

**Authors:** Alexander J Hull, Magda L Atilano, Kerri J. Kinghorn

## Abstract

Gaucher disease (GD), the most common lysosomal storage disorder, is caused by bi-allelic mutations in the *GBA1* gene. Variants in *GBA1* also represent the most frequent genetic risk factor for Parkinson’s disease (PD). Although GD and PD are clinically distinct disorders, they share key pathological features, including lysosomal dysfunction, mitochondrial stress, and redox imbalance. While PD has traditionally been studied in the context of neuronal decline, the contribution of non-neuronal organ systems remains poorly understood. Here, we demonstrate that progressive renal dysfunction is a central, disease-modifying feature in *Drosophila* models of GD and PD. We show that *Drosophila* lacking either the main fly orthologue of *GBA1*, *Gba1b*, or the mitophagy regulator *Parkin*, exhibit age-dependent degeneration of the renal system. This includes disorganisation of the Malpighian tubules, impaired nephrocyte function, redox imbalance, and lipid accumulation. These renal defects contribute to systemic physiological decline, including water retention, ionic hypersensitivity, and exacerbation of neurodegenerative phenotypes. Importantly, we identify redox dyshomeostasis, rather than classical oxidative stress, as a central pathogenic driver, marked by paradoxical sensitivity to both oxidative and reductive interventions. Notably, treatment with the mTOR inhibitor rapamycin selectively restores renal structure and function in *Gba1b* mutants, but not in *Parkin* mutants, revealing mechanistic divergence between lysosomal and mitochondrial stress. These findings uncover redox imbalance as a biomarker of renal vulnerability and establish the renal system as a critical, potentially disease-modifying organ in the systemic progression of GD and PD.

## Introduction

Bi-allelic mutations in the *GBA1* gene, which encodes the lysosomal enzyme glucocerebrosidase (GCase), are the primary cause of the lysosomal storage disorder Gaucher disease (GD). Moreover, heterozygous *GBA1* mutations represent a significant genetic risk factor for Parkinson’s disease (PD) (Migdalska-Richards & Schapira, 2016). GCase plays a crucial role in lysosomal homeostasis by facilitating the breakdown of glucosylceramide (GluCer) into ceramide and glucose (Boer et al., 2020). Loss of normal GCase function in GD leads to the accumulation of lipid-laden Gaucher cells in multiple organs, primarily in the liver, spleen, and bone marrow. These cells may also accumulate in other tissues, including the lymphatic system, lungs, skin, eyes, kidneys, heart, and, in rare cases, the nervous system (Cox, 2010). While the neurological and haematological manifestations of GD are well-documented, the impact of *GBA1* mutations on renal function remains an underexplored but emerging area of interest. Furthermore, the potential involvement of the renal system in PD is also becoming increasingly recognised (Meléndez-Flores & Estrada-Bellman, 2021).

Gaucher macrophages have been identified within endothelial and interstitial regions of the kidney, where they may contribute to severe proteinuria. However, renal dysfunction remains a rare symptom in GD patients (Smith et al., 1978; Chander et al., 1979; Siegel et al., 1981; Halevi et al., 1993). GluCer accumulation has also been observed in the glomeruli of patients with diabetic nephropathy and polycystic kidney disease (PKD), as well as in rodent models (Shayman et al., 1991; Zador et al., 1993; Deshmukh et al., 1994; Natoli et al., 2010). Furthermore, treatment with GluCer synthase inhibitors has been shown to ameliorate glomerular degeneration in PKD, and clinical trials are underway, highlighting the critical role of GluCer regulation in maintaining renal health (Natoli et al., 2010; Shayman et al., 2016).

Loss of function mutations in *LIMP2* (also known as *SCARB2*), a sorting receptor that specifically directs GCase to the lysosome, have been identified in multiple families as the cause of progressive action myoclonus and renal failure (AMRF) (Berkovic et al., 2008; Balreira et al., 2008; Chaves et al., 2011; Hopfner et al., 2011). In addition to myoclonus, AMRF is marked by proteinuria, weight gain, and fluid retention leading to oedema. The mechanism underlying *LIMP2*-associated AMRF is likely distinct from GD, as GluCer-filled macrophages are not observed in tissue samples from AMRF patients. Although not directly implicated in renal pathology, AMRF patients and *Limp2* knockout mice show a several fold increase in circulating levels of glucosylsphingosine, a water soluble deacylated form of GluCer that can be excreted in urine. This increase is also seen in GD (Dekker et al., 2011; Gaspar et al., 2014). Furthermore, renal failure is a major complication in Fabry disease, another lysosomal storage disorder characterised by the accumulation of the complex glycosphingolipid globotriaosylceramide (GB3) (Nakao et al., 2003).

An emerging link between PD and kidney dysfunction is becoming increasingly evident. Large cohort studies involving patients with nephrotic syndrome, end-stage renal disease (ESRD), and chronic kidney disease (CKD) have consistently reported a significantly higher incidence of PD or parkinsonism (Huang et al., 2018; Wang et al., 2014; Melendez-Flores, 2021). However, it remains unclear whether renal and neurological diseases directly influence one another or whether they share a common upstream pathological mechanism. Evidence from large datasets, including South Korean populations and UK Biobank samples, suggests that a reduced glomerular filtration rate (GFR) and elevated proteinuria are associated with an increased risk of developing PD (Nam et al., 2019; Peng et al., 2024). In individuals with PD, an inverse correlation has been observed between GFR and serum levels of neurofilament light chain (NfL), a biomarker of neuronal damage, as well as between GFR and cognitive decline (Qu et al., 2023). These findings imply that kidney dysfunction is closely linked to the severity of PD. Notably, both PD and various forms of kidney disease are characterised by elevated oxidative stress, suggesting that shared pathophysiological mechanisms may underlie this association (Pizzino et al., 2017).

Fruit flies are increasingly used as models for renal disorders. First, they serve as a kidney stone model, as xanthic/uric acid crystals can form in the *Drosophila* Malpighian tubules (MTs) (Dow & Romero, 2010). Second, they are used to study podocyte biology, as the slit diaphragm structure of mammalian podocytes is conserved in fly nephrocytes, a large class of cells adjacent to the heart tube that filter the haemolymph through fluid-phase endocytosis (Weavers et al., 2009). Water retention can result from the loss of filtration efficiency in the MTs, caused by the breakdown of occluding junctions between epithelial cells or impaired renewal of differentiated renal epithelial cells (Martinez-Corrales et al., 2016; Denholm et al., 2010; Dornan et al., 2023).

Using our previously characterized GD *Drosophila* model lacking *Gba1b*, the primary fly orthologue of *GBA1*, we demonstrate progressive degeneration of the renal system, loss of organismal water homeostasis and ionic hypersensitivity. Specifically, *Gba1b^-/-^* flies exhibit reduced glomerular filtration capacity in pericardial nephrocytes, along with widespread cellular degeneration in the MTs. Furthermore, renal system dysfunction negatively impacts both organismal and neuronal health in *Gba1b^-/-^*flies, including autophagic-lysosomal status in the brain. Additionally, we show that the loss of Parkin, a familial PD-linked E3 ubiquitin ligase, similarly leads to renal defects. Notably, renal dysfunction in GD flies can be alleviated through rapamycin administration, highlighting a potential therapeutic avenue for lysosomal and neurodegenerative diseases linked to GBA1 dysfunction.

## Materials and Methods

### Fly stocks and rearing

Flies were reared at 25°C and 50% relative humidity in a 12:12LD cycle on SYA Food (1.5% agar, 5% sucrose, 10% dried yeast [Leiber Bierhefe], 0.3% nipagin, 0.3% propionic acid). Experimental flies were raised at a defined density, mated for 1 day and females were subsequently maintained at a density of 15 flies per vials and changed to fresh food every 2-3 days. The following fly stocks were acquired from the Bloomington *Drosophila* Stock Center: *Gba1b*^CRIMIC^ (BL#78943), *park^1^* (BL#34747), C724-Gal4 (BL#99567), c42-Gal4 (BL#30825), tub-Gal4 (BL#5138), UAS-*CD8RFP* (BL#27392), UAS-mcherry.nls (BL#38424), UAS-mitoGFP (BL#8442), UAS-yfpPTS (BL#64247), *klf15* (BL#18979), UAS-*Sod1* (BL#24754), UAS-*Sod2* (BL#24494), UAS-*CatA* (BL#24621), UAS-mitoroGFP2.grx1 (BL#67664), UAS-*cytoroGFP2*.grx1 (BL#67662), UAS-*Park* (BL#51651). *Gba1b^-/-^* flies were previously constructed by homologous recombination (Kinghorn et al., 2016). UAS-*Nrf2* and UAS-*Keap1* were gifts from Dirk Bohmann (Sykiotis & Bohmann, 2008) and UAS-mitoQC was a gift from Alex Whitworth (University of Cambridge; Lee et al., 2016).

### Water weight assays

Flies were frozen in liquid nitrogen and immediately weighed on an AT-201 Mettler Toledo Analytical Balance. Flies were then dried out for 24 hrs in an 80°C oven and re-weighed. The total water volume was calculated as the weight lost from drying.

### Hemolymph extraction

Groups of 15 flies were beheaded, placed upside-down in a truncated pipette tip in a 1.5ml Eppendorf and centrifuged for 10 minutes at 1500 rpm at 4°C. The haemolymph was then flash-frozen in liquid nitrogen.

### Drinking assays

Assay 1: Groups of 15 female flies were placed in vials containing dried fly food and a 1.5 mL Eppendorf tube filled with water. The Eppendorf tubes were weighed before and after a 24-hr period. The difference in weight, adjusted for evaporation using a control tube lacking a fly, was calculated and normalised per fly. Assay 2: Cohorts of flies were starved for 5 hours before being provided with a water source for 1 hour. To estimate water consumption, the mean weight of water in the starved cohort was subtracted from the weight of flies given access to water.

### Cold-stress assays

>75 flies of each genotype were placed in empty glass tubes and uniformly submerged in an ice bucket for 10 minutes. To assess chill-coma recovery, flies were filmed and considered recovered once they stood upright and began walking. Survival was scored periodically over a 3-day period.

### Lifespan

Lifespan assays were conducted as previously described (Atilano et al., 2023). Eggs were collected over a 24-hour period on grape juice agar plates and then pipetted onto SYA food at a uniform density. Female flies were split into groups of 15 in vials, transferred to fresh food every 2–3 days, and monitored for survival.

### Ionic stress and natriuresis assays

Experimental flies were raised on SYA food as described above for three weeks before being transferred to food containing 4% NaCl or KCl, with or without a water source provided by 1% agar in a pipette tip (Van Dam et al., 2020). For dehydration assays, flies were raised without a water source on active dry yeast (*Saf-Levure*).

### FRUMS (Fluorescent Renal Uptake and Metabolite Secretion) assay

A volume of 23 nL of 2.5% (w/v) Brilliant Blue FCF-E133 (Florio Colori) in ddH₂O was injected into the haemolymph using a Drummond Nanoject II Microinjector. Blue flies were then scored manually every hour for qualitative clearance of dye from the hemolymph and subsequently from the MT and gut.

### Tissue preparation and immunostaining

MTs and brains were dissected in ice-cold PBS and immediately fixed in 4% PFA for 20 minutes. The tissues were then washed three times for 20 minutes each in PBS-T (0.5% Triton-X), followed by a 30-minute incubation in blocking solution (5% Normal Goat Serum in PBS-T). Primary antibody incubation was performed overnight at 4°C with gentle nutation. Tissues were then washed three times for 20 minutes in PBS-T and incubated with secondary antibodies in blocking solution for 2 hours at room temperature. This was followed by three additional 20-minute washes in PBS-T before mounting in Vectashield Mounting Medium with DAPI (Vector Laboratories). Primary antibodies used were mouse anti-cora C615.15 (DSHB,1:200), rabbit anti-Ref(2)P (ab178440, 1:200), mouse anti-FK2 (Sigma 04263, 1: 800). The secondary antibodies used were goat anti-mouse Alexa488 (A11001) and goat-rabbit Alexa568 (A11036), both at a dilution of 1:250. For BODIPY experiments, 20 µM BODIPY^TM^ 505/515 was incubated 1:100 in addition to the primary antibody.

Tissues with genetically encoded fluorescent markers (YFP-PTS, mito-Grx1-roGFP2 and cyto-Grx1-roGFP2) were dissected in 20 mM NEM in PBS, fixed in 4% PFA for 20 minutes, washed for 20 minutes in PBS-T, and mounted in PBS.

### Dextran uptake assay

Fly abdomens were dissected from 28-day-old flies and incubated in 0.33 mg/mL Texas Red 10,000 MW dextran (Thermo Fisher) in PBS for 5 minutes. The abdomens were then washed twice in PBS, fixed in 4% PFA for 20 minutes, and washed for 20 minutes in PBS-T. Finally, the samples were mounted in VECTASHIELD mounting medium with DAPI and imaged on a Zeiss 700 confocal microscope.

### Live imaging

Tissues were dissected at room temperature and transferred to 50 nM Lysotracker^TM^ Red DND-99 (Thermo Fisher), Mitotracker^TM^ Red CMH_2_XROS, or DHE (Thermo Fisher) in PBS, followed by immediate imaging at 20x magnification using a Zeiss 700 or 880 confocal microscope. For mitoQC experiments, MTs were dissected in cold PBS and immediately imaged. For lipid peroxidation analysis, MTs were dissected in PBS and incubated in 20 µM BODIPY^TM^ 581/591 in Schneider’s medium for 30 minutes. After two PBS washes, tissues were mounted in PBS and immediately imaged

### Reduced glutathione (GSH) assay

GSH levels were measured using a Glutathione Assay Kit (Cayman Chemicals) according to the manufacturer’s instructions. Briefly, groups of 10 flies were homogenised with glass beads (Sigma) in MES buffer using a Precellys Ribolyser. The homogenate was then centrifuged at 10000 g for 10 minutes and the resulting supernatant was transferred to a new tube. Samples were deproteinated in 50 mg/ml metaphosphoric acid and basified with 1/20 4M triethanolamine. A 50 µl aliquot of each sample was loaded into Greiner transparent flat-bottom 96-well plates (Cellstar) containing 150 µl enzyme buffer. Absorbance was measured at 405 nm at 5-minute intervals for 30 minutes using a Tecan Infinite M200 plate reader. GSH values were calculated from the slope of a standard curve generated using known Glutathione Persulfide (GSSG) concentrations.

### NAD/NADH assay

NAD/NADH quantification was performed using a NAD+/NADH Quantification Colorimetric Kit (Biovision) according to the manufacturer’s instructions. Groups of 10 flies were flash-frozen in liquid nitrogen and homogenized in 300 µL of extraction buffer. Samples were deproteinated by centrifugation at 10000 rpm for 10 minutes at 4°C using Amicron Ultra 0.5 mL 10,000 MW filters. The samples were then divided into NAD and NADH groups, where NAD was decomposed by incubation at 60°C for 1 hour. Following this, samples were incubated in NAD cycling buffer and developer, and absorbance readings were taken at 450 nm using a Tecan Infinite M200 plate reader.

### Statistical analyses

Data was analysed in Graphpad Prism 8.0.2 and the statistical tests used are described in the figure legends. p<0.05 was considered statistically significant in all cases. Log-rank test on lifespan data were performed in Microsoft Excel (available at http://piperlab.org/resources/).

## Results

### Flies lacking *Gba1b* display age-dependent MT degeneration

Since both sphingolipid accumulation and PD are linked to renal disease, we aimed to assess the renal system in a previously characterised GD *Drosophila* model, lacking the main fly orthologue of *GBA1*, known as *Gba1b* (Kinghorn et al., 2016). Fly MTs consist of a tubulated epithelial sheet composed of principal cells (PCs) and stellate cells (SCs) surrounding a lumen. To investigate structural integrity, we stained the MTs with coracle (Cora), a core component of the septate junction (SJ), a specialised cell-cell junction which is essential for barrier function and transcellular water flow (Dornan et al., 2023). In control fly MTs, Cora marked clearly defined cell vertices throughout the tubule. However, in *Gba1b^-/-^*fly MTs, the number of well-defined SJ vertices was significantly reduced, and Cora was frequently mislocalised throughout the cytoplasm (Figure 1A). Notably, these structural abnormalities were absent in day 1 *Gba1b^-/-^* fly MTs, indicating that the loss of tissue architecture is progressive.

**Figure 1.**
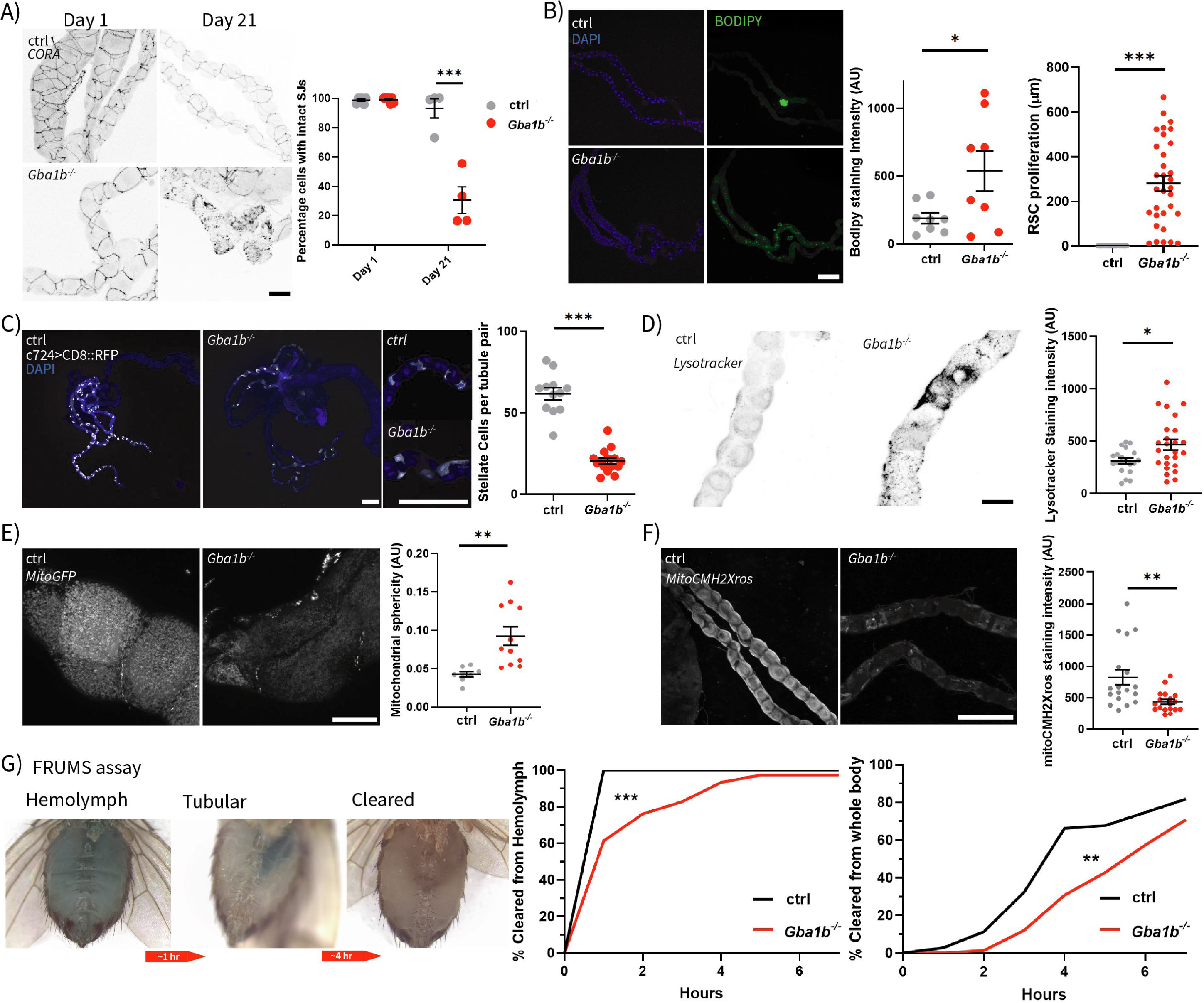
*Gba1b^-/-^*flies show a progressive deterioration of MT morphology and function. A) Representative images (10× magnification) and quantification of Coracle (CORA; grayscale) immunostaining at day 1 and 21, illustrating the progressive loss of tubule junctional integrity in aged *Gba1b*^-/-^ flies compared to controls (ctrl) (two-way ANOVA with Fisher’s LSD multiple comparisons, Day 1 p=0.9547, Day 21 ***p<0.0001), Scale-bar = 100µm. B) Representative images of tubules stained with BODIPY^TM^ (green) and DAPI (blue) and quantification showing increased neutral lipid and RSC proliferation in 3-week-old *Gba1b^-/-^* fly tubules compared to ctrl (BODIPY^TM^ quantification, unpaired t-test, *p=0.0374; DAPI RSC proliferation, unpaired t-test, ***p<0.0001)). Scale-bar = 200µm. C) Representative images and quantification of stellate cell (SC) number in 3-week-old ctrl and *Gba1b^-/-^* flies, visualised using c724>CD8::RFP, show a reduction in SCs in *Gba1b^-/-^* flies compared to ctrl (unpaired t-test, ***p<0.0001). Scale-bar = 250µm. D) Representative images and quantification of lysosomal staining intensity in MTs, visualised with LysoTracker^TM^ Red, showing increased lysosomal enrichment in 3-week-old *Gba1b*^-/-^ flies compared to controls (unpaired t-test, *p=0.0144). Scale-bar = 50µm. E) Representative images of the mitochondrial morphology in the tubules of ctrl and *Gba1b*^-/-^ flies, visualised using tub>mitoGFP and quantified for sphericity with the Mitochondrial Analyser plugin. The analysis reveals that mitochondria in *Gba1b*^-/-^ fly tubules are less fused and exhibit a more punctate morphology compared to ctrl (ctrl vs *Gba1b^-/-^*, **p=0.0032, unpaired t-test). Scale-bar = 25µm. F) Representative images and quantification of MitoTracker™ Red CM-H₂XRos staining in renal tubules, showing reduced mitochondrial membrane potential in 3-week-old *Gba1b^-/-^* flies compared to ctrl (unpaired t-test, **p=0.0036). Scale-bar = 200µm. G) Example images of a FRUMS assay, showing the clearance of a systemically injected dye from the hemolymph, the tubule and the whole fly. *Gba1b^-/-^* flies display reduced filtration and removal of the dye from the haemolymph and whole body compared to ctrl (ctrl vs *Gba1b^-/-^* cleared from the hemolymph, ***p<0.0001; ctrl vs *Gba1b^-/-^* cleared from whole body; **p=0.0012, log-rank tests).

Since lipid aggregation is a hallmark of metabolic disorders, we stained MTs with the lipid marker BODIPY, which revealed a significant increase in luminal lipids in *Gba1b^-/-^* flies compared to controls (Figure 1B). Moreover, counter-staining with the nuclear marker DAPI identified a significant increase in small nuclei proliferating from the ureter, presumptively corresponding to renal stem cells (RSCs), extending into the absorptive zone of the MT in *Gba1b^-/-^* flies compared to age-matched control MTs. RSCs are known to proliferate in response to stress and inflammation (Singh et al., 2007), resulting in decreased water flux (Martinez-Corrales et al., 2019).

Transcellular water flux principally occurs through SCs, and given the observed loss of normal tissue architecture, we quantified SCs in *Gba1b^-/-^* fly MTs through expression of the membrane-localised CD8:RFP using the SC-specific driver C724-Gal4. We observed a significant reduction in SC numbers in *Gba1b^-/-^* fly MTs compared to those seen in control flies, alongside a loss of their characteristic star or bar morphology (Figure 1C).

The gut and brain of *Gba1b**^-/-^*** flies, similar to macrophages in GD patients, are characterised by enlarged lysosomes (Kinghorn et al., 2016, Atilano et al 2023). Consistent with this, staining of MTs with LysoTracker^TM^, an acidic vesicle dye, revealed a significant increase in fluorescence intensity in *Gba1b^-/-^* fly MTs, consistent with pronounced lysosomal expansion in a subset of renal cells (Figure 1D).

As MTs play a critical role in redox balance, with mitochondrial activity in PCs being pivotal to physiological function, we examined mitochondrial morphology using mito-GFP expression. In control PCs, mitochondria appeared fused and tubulated, whereas in *Gba1b^-**/-**^* fly MTs, they were predominantly punctate, indicative of defective mitochondrial fusion-fission homeostasis (Figure 1E). We next stained with MitoTracker^TM^ CM-H2XRos, a mitochondrial-specific fluorescent probe used to measure reactive oxygen species (ROS). This demonstrated reduced staining intensity in the MTs of *Gba1b^-/-^* flies. This is likely consistent with a decrease in mitochondrial ROS production, due to impaired mitochondrial membrane potential and dysfunction of the electron transport chain (ETC) (Figure 1F). In keeping with this, NADH, but not NAD+, was decreased in whole *Gba1b^-/-^* flies, suggesting impaired mitochondrial energy metabolism (Figure S1A).

Finally, to assess MT physiological function, we employed the FRUMs assay, in which blue food dye is injected into the hemolymph, and its clearance into the tubules and subsequent excretion serve as a proxy for water homeostasis. *Gba1b^-/-^* flies exhibited increased retention of blue dye in both the hemolymph and subsequently the tubules, indicating impaired MT-mediated hemolymph clearance (Figure 1G).

### Ion and water homeostasis is dysregulated in *Gba1b^-/-^* flies

To investigate the impact of these defects in MT morphology and function on water homeostasis, we measured water weight in aged *Gba1b^-/-^*flies. Our analysis revealed an increase in both total water weight and percentage body water weight (Figure 2A), partially attributable to an elevated hemolymph volume in *Gba1b^-/-^* flies compared to controls (Figure 2B). Notably, this increase was not due to heightened water intake, as *Gba1b^-/-^* flies exhibited either reduced or unchanged drinking behaviour across multiple assays (Figure S2A-B).

**Figure 2.**
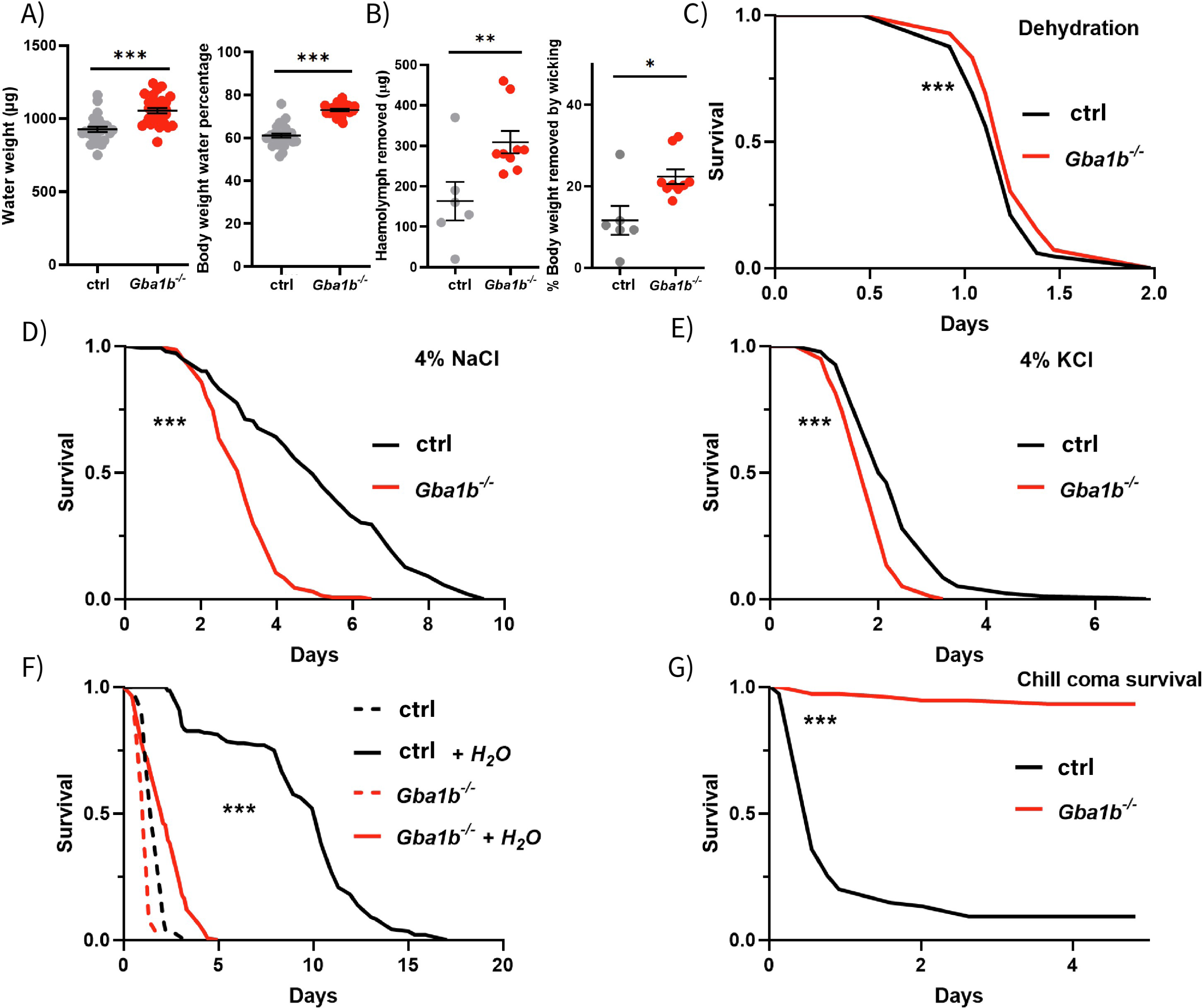
*Gba1b^-/-^* flies exhibit impaired ionic homeostasis. A) 3-week-old *Gba1b^-/-^* flies display a greater body water weight and body weight water percentage compared to ctrl flies (unpaired t-test, ***p<0.0001; ***p<0.0001). B) Hemolymph water weight is also increased in *Gba1b^-/-^*flies compared to ctrl by hemolymph removal and by wicking (unpaired t-test, water removed; **p=0.0132; percentage body weight removed *p=0.0112). C) Dehydration assay showing that 3-week-old *Gba1b^-/-^*flies are more resistant to desiccated food than age-matched ctrl (log-rank test, ***p=0.0013). D) 3-week-old *Gba1b^-/-^* flies are more sensitive to a diet supplemented with 4% NaCl and display reduced survival compared to ctrl (log-rank test, ***p<0.0001). E) The survival of *Gba1b^-/-^* flies is reduced compared to ctrl on a diet supplemented with 4% KCl (log-rank test, ***p<0.0001). F) A natriuresis assay showing the survival curves for *Gba1b^-/-^* and ctrl flies on diet supplemented with 4% NaCl with or without addition of H_2_O source (log-Rank test, ctrl vs *Gba1b^-/-^* flies, ***p < 0.0001; ctrl vs ctrl + H_2_O, ***p<0.0001; *Gba1b^-/-^* vs *Gba1b^-/-^* + H_2_O, ***p < 0.0001). G) 3-week-old ctrl flies are strongly sensitive to cold shock compared to *Gba1b^-/-^* flies (log-rank test, ***p < 0.0001).

To determine the physiological consequences of this water retention, we subjected *Gba1b^-/-^* flies to a desiccative stress assay and observed that they exhibited a small but significant dehydration resistance compared to controls (Figure 2C). *Gba1b^-/-^* flies also retained a higher level of water throughout the course of the dehydration assay (Figure S2C). As MTs play a crucial role in regulating ion homeostasis, we subjected *Gba1b^-/-^*flies to ionic stress assays using food supplemented with 4% NaCl or KCl. We observed that *Gba1b* mutants were highly sensitive to both ionic stress conditions (Figure 2D, E). To determine whether the lethality observed in these assays was due to desiccative stress from water loss accompanying excess salt efflux through the MTs, or direct cellular ionic stress, we hypothesised that providing a water source should enable flies with functional MTs to maintain homeostasis under ionic stress conditions. To test this, we measured both water levels and lifespan under ionic stress while supplementing a water source via 1% agar-filled pipette tip (Figure 2F, S2D). Water supplementation led to a substantial lifespan extension in control flies exposed to 4% NaCl, with a smaller yet significant extension in *Gba1b^-/-^*flies (Figure 2F). NaCl exposure led to a reduction in water weight, which was rescued by water supplementation in control but not *Gba1b^-/-^* flies (Figure S2D). These observations suggest that the early mortality in control flies under ionic stress primarily results from dehydration rather than direct ionic toxicity. On the other hand, although dehydration contributes to *Gba1b^-/-^* mortality, a significant component of their susceptibility to ionic stress is likely due to defective natriuresis, leading to impaired ion homeostasis.

In *Drosophila*, cold tolerance is closely linked to altered ion homeostasis. To investigate this, we subjected aged control and *Gba1b^-/-^*flies to a chill-coma stress assay and found that flies lacking G*ba1b* exhibited a remarkable resistance to chill-induced damage, whereas control flies experienced rapid mortality within 24 hours (Figure 2G).

### Hemolymph clearance and nephrocyte function is diminished in *Gba1b^-/-^*flies

Podocyte-like nephrocytes filter small macromolecules through a size-excluding slit-diaphragm system, after which they are then taken up non-selectively through fluid-phase endocytosis and degraded in the lysosome (Weavers et al., 2019). To assess endocytic function, we incubated dissected pericardial nephrocytes with Texas-red dextran to visualise early endosomes. At 4 weeks of age, *Gba1b^-/-^* flies exhibited a population of endosome-negative nephrocytes, indicating a loss of endosomal activity, and consequently, hemolymph filtration (Figure 3A). Consistent with reduced filtration, protein concentration analysis using a BCA assay on extracted hemolymph revealed a significant increase in circulating soluble protein in *Gba1b^-/-^* flies relative to controls (Figure 3B). These findings suggest that defects in both MT function and glomerular filtration contribute to increased circulating waste, potentially leading to secondary toxic consequences.

**Figure 3.**
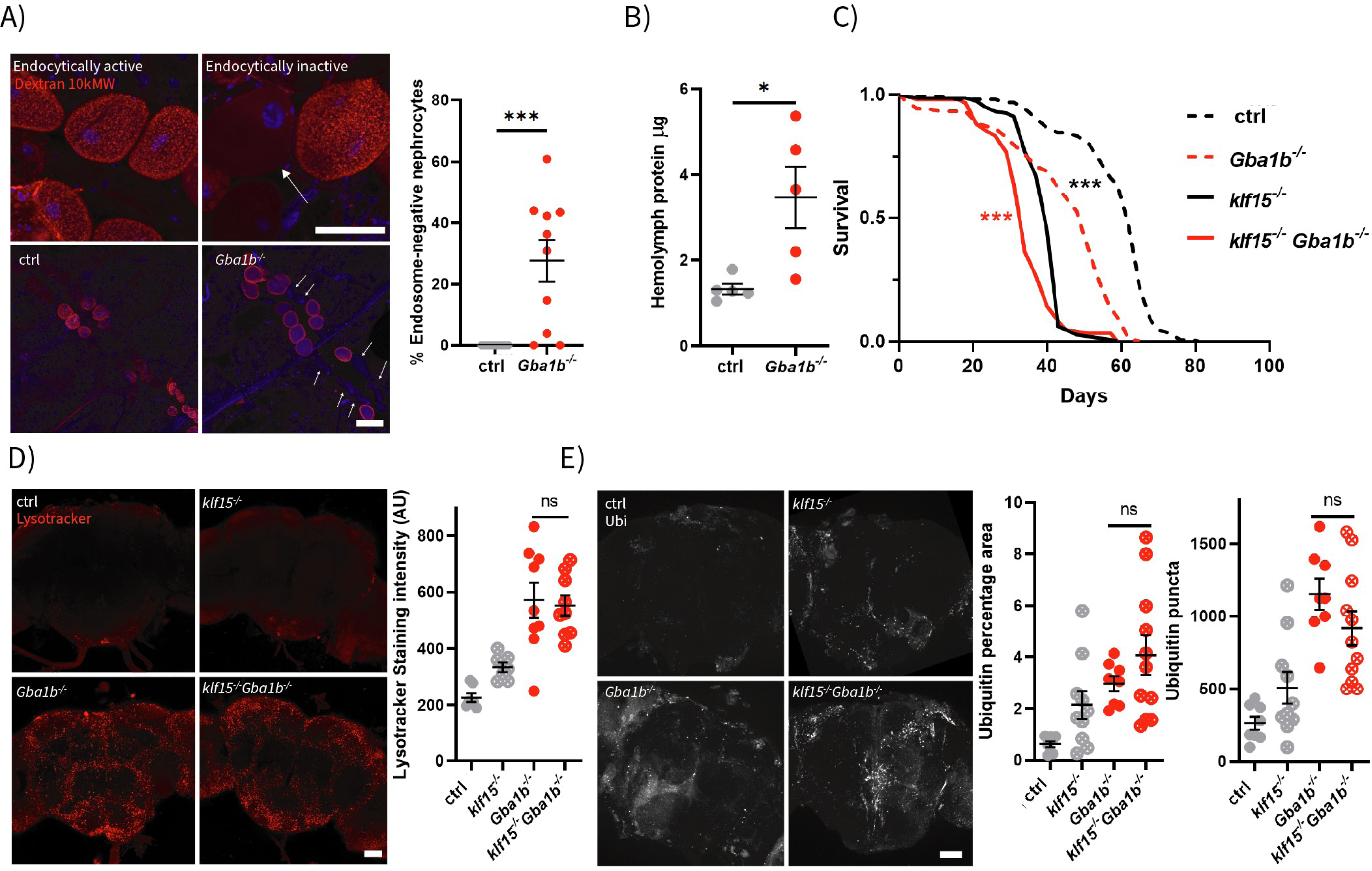
Aged *Gba1b^-/-^* flies display diminished hemolymph clearance and nephrocyte function. A). Representative 63x and 10x images and quantification of nephrocytes from ctrl and *Gba1b^-/-^*flies at 4 weeks of age following uptake of Dextran 10,000 MWA. Dextran uptake (Red) represents endocytic activity and DAPI (Blue) labels nuclei. In *Gba1b^-/-^*fly nephrocytes, a subset of cells lack Dextran (arrow) (unpaired t-test, ***p < 0.0001). Scale bar for 10x and 63x = 100 µm. B) The soluble protein content of the hemolymph of *Gba1b^-/-^* flies is higher than in ctrl flies as assessed by a BCA assay (unpaired t-test, \**p*=0.0180), indicating impaired filtration. C). Lifespan survival curves of *klf15^-/-^*; *Gba1b^-/-^* double mutants and ctrl. Loss of *klf15* significantly reduces lifespan in both ctrl and *Gba1b^-/-^* flies (log-rank test, *Gba1b^-/-^* vs *klf15^-/-^; Gba1b^-/-^*, \*\*\**p* < 0.0001). D) Representative 20× images of LysoTracker^TM^ (Red)-stained brains from *klf15^-/-^; Gba1b^-/-^* flies and ctrl. Quantification of lysosomal area and puncta number shows no significant difference between *Gba1b^-/-^* and *klf15^-/-^; Gba1b^-/-^* genotypes (one-way ANOVA with Tukey’s test, *p*=0.9830). Scale bar = 50 µm. E) Representative 20x images of FK2/polyubiquitin (grayscale) immunostaining in brains from *klf15^-/-^; Gba1b^-/-^* flies and ctrl. Quantification of polyubiquitin puncta and area reveals no significant difference between *Gba1b^-/-^* and *klf15^-/-^; Gba1b^-/-^* genotypes (one-way ANOVA with Tukey’s test: FK2 puncta *p*=0.4015; FK2 area *p*=0.5228). Scale bar = 50 µm.

To determine the contribution of nephrocyte dysfunction to disease pathology in *Gba1b* loss of function, we impaired nephrocyte maturation using a *Klf15* LOF mutant, which is exclusively expressed in nephrocytes and required for their differentiation (Ivy et al., 2018). Loss of *Klf15* significantly shortened the lifespan of both *Gba1b^-/-^*and control flies, demonstrating that nephrocyte dysfunction is deleterious to both healthy flies and those lacking GCase activity (Figure 3C). To determine the consequence of abnormal nephrocyte function on brain health, we measured lysosomal area in the brain with LysoTracker^TM^, or stained for polyubiquitin. *Klf15* LOF had no significant effect on lysosomal or polyubiquitin area relative to healthy controls, nor an additive effect on the increased lysosomal volume (Figure 3D) and polyubiquitin accumulation (Figure 3E) seen in *Gba1b^-/-^* fly brains. This suggests that loss of nephrocyte viability itself does not exacerbate brain pathology.

### *Gba1b^-/-^* MTs display loss of redox homeostasis capacity

The renal system is key to organismal redox homeostasis, a process thought to play a key role in the pathogenic mechanisms of GD and PD progression (Kishi et al., 2024). Glutathione (GSH), the most abundant antioxidant in *Drosophila*, plays a crucial role in redox homeostasis by reducing hydrogen peroxide (H₂O₂) and lipid hydroperoxides (ROOH) via GSH peroxidases. To evaluate redox balance, we measured total GSH levels in whole flies from both control and *Gba1b^-/-^* groups. Our results revealed a significant decrease in GSH in *Gba1b^-/-^*flies, indicating a compromised ability to maintain redox equilibrium (Figure 4A). To further evaluate the glutathione redox potential (E_GSH_) in MTs, we utilised the redox-sensitive green, fluorescent biosensor Grx1-roGFP2, targeted to both the mitochondria and cytosol (Albrecht et al., 2011). Our findings revealed a shift toward a more reduced glutathione state in the mitochondria of *Gba1b^-/-^*fly tubules compared to controls (Figure 4B). Interestingly, while cytosolic glutathione typically maintains a less oxidised GSH/GSSG ratio than mitochondria, it was paradoxically more oxidised in *Gba1b^-/-^* tubules relative to controls (Figure 4C). This indicates that despite a potential reduction in mitochondrial ROS production, the tubules fail to effectively regulate basal ROS levels. These abnormalities may contribute to the impaired physiological function of the MTs.

**Figure 4.**
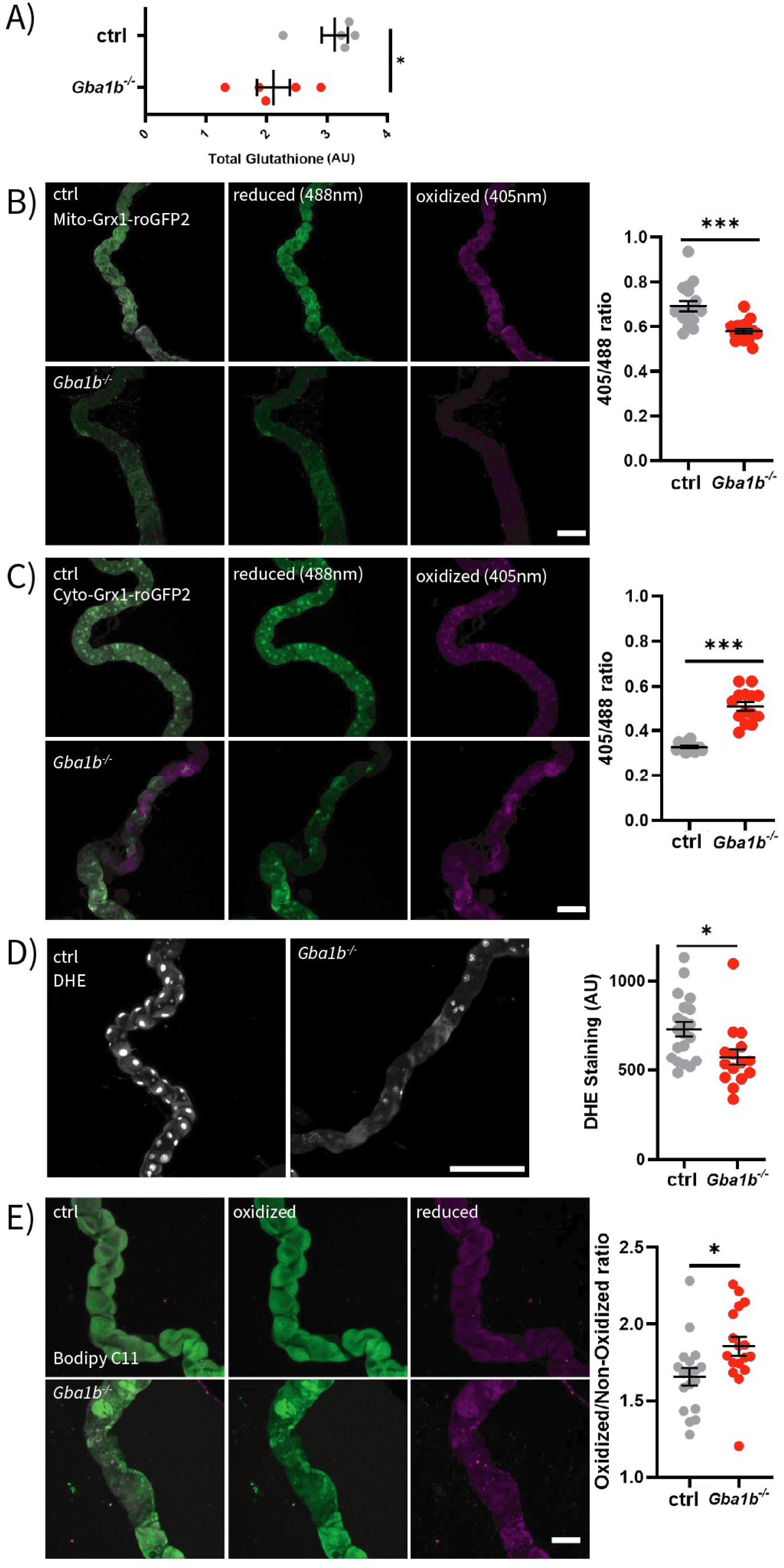
Disruption of redox homeostasis in *Gba1b^⁻/⁻^* flies drives disease-associated phenotypes. A) Whole-fly total glutathione (GSH) abundance is significantly decreased in 3-week-old *Gba1b^-/-^* flies compared to ctrl flies (unpaired t-test, \**p*=0.019). B) Representative 20× images and quantification of the 405 nm (oxidised)/488 nm (reduced) excitation ratio using the mitochondrial-targeted Grx1-roGFP2 redox biosensor in *tub>mito-Grx1-roGFP2* control and *Gba1b^-/-^*flies. *Gba1b^⁻/⁻^* fly tubules exhibt a significantly lower 405/488 ratio, indicating a shift toward a more reduced glutathione redox state (unpaired t-test, \*\*\**p*<0.0001). Reduced signal is shown in magenta, oxidised in green. Scale bar = 100 µm. C) Example 20× images and quantification of the 405 nm (oxidised)/488 nm (reduced) ratio in tub>*cyto-Grx1-roGFP2* ctrl and *Gba1b^-/-^*fly MTs. Quantification shows a significant increase in oxidation in *Gba1b^-/-^* tubules (unpaired *t*-test, \*\*\**p* < 0.0001). Scale bar = 100 μm. D) Representative 20× images and quantification of dihydroethidium (DHE) fluorescence intensity in control and *Gba1b^-/-^* MTs. DHE staining is significantly reduced in *Gba1b^-/-^* fly tubules compared to ctrl (unpaired t-test, **\***p = 0.0151). Scale bar = 200 µm. E) Representative images and quantification of lipid peroxidation in MTs stained with the polyunsaturated lipid peroxidation sensor BODIPY^TM^ 581/591. Oxidised lipids are shown in green (excitation: 488 nm), and reduced lipids in magenta (excitation: 568 nm). *Gba1b^-/-^* mutant MTs exhibit significantly elevated oxidised/reduced lipid ratios compared to ctrl (unpaired t-test, *p =0.0242). Scale bar = 100 µm.

We then incubated MTs with the nuclear-localised superoxide indicator dihydroethidium (DHE), revealing a significant decrease in superoxide levels in *Gba1b^-/-^* tubules (Figure 4D). Since glutathione primarily detoxifies H₂O₂ and ROOH, rather than superoxide, this reduction could indicate lower mitochondrial ROS production or altered superoxide metabolism (Winterbourn, 2016).

To investigate the role of lipids in redox dyshomeostasis, we measured lipid peroxidation in MTs using the polyunsaturated lipid peroxidation marker BODIPY^TM^ 581/591 C11. *Gba1b^-/-^* fly tubules exhibited higher lipid peroxidation ratios than controls, suggesting increased oxidative damage and potential deficiencies in antioxidant defences or lipid turnover mechanisms (Figure 4E). We conclude that *Gba1b^-/-^* flies, particularly their MTs, have a diminished capacity to regulate redox balance.

To determine the endogenous antioxidant defence in the tubules, we profiled peroxisomal morphology using the marker YFP-PTS. We found a significant decrease in peroxisome density and area in *Gba1b^-/-^* fly tubules, potentially indicating defects in peroxisome maturation and function (Figure S3A). We further investigated peroxisomal and cytosolic antioxidant defences by immunostaining for superoxide dismutase 1 (Sod1) and catalase (CatA). Sod1 showed a significant decrease in staining intensity and a transition from a compartmentalised to diffuse localisation in *Gba1b^-/-^* fly tubules (Figure S3B). Taken together, these findings indicate that the MTs of *Gba1b^-/-^* flies exhibit redox imbalance characterised by mitochondrial, peroxisomal, and lipid oxidative stress. Moreover, the inability to effectively regulate antioxidant responses may have broader implications for physiological functions across multiple tissues in *Gba1b* deficiency.

### Induction of antioxidant effector Nrf2 worsens renal and neurological phenotypes in *Gba1b^-/-^* flies

To investigate the contribution of redox dyshomeostasis to *Gba1b^-/-^*fly disease phenotypes, we overexpressed the antioxidant regulator Nrf2/CnC in PCs, a manipulation previously shown to prematurely impair renal function (Burbridge et al., 2021). Consistent with this, whilst Nrf2 overexpression resulted in a modest but statistically significant lifespan reduction in control flies, its overexpression in *Gba1b^-/-^* flies led to a dramatic lifespan shortening (Figure 5A). This suggests that *Gba1b^-/-^* flies have an increased susceptibility to antioxidant-mediated tubular dysfunction. In addition, we observed that Nrf2 overexpression in *Gba1b^-/-^* flies resulted in an increase in renal stem cell proliferation at the ureter (Figure 5B). However, despite these effects, there was a significant increase in water weight in controls overexpressing Nrf2, but not in *Gba1b^-/-^* flies (Figure 5C). Collectively, these findings indicate that *Gba1b^-/-^* flies exhibit disproportionate negative renal and systemic responses to activation of the antioxidant response. We next tested the consequence of Nrf2 overexpression in PCs on neuronal health in *Gba1b* loss of function. This intervention led to a significant increase in lysosomal puncta number, as assessed by LysoTracker^TM^ staining (Figure 5D), and exacerbated protein dyshomeostasis, as indicated by polyubiquitin accumulation and increased levels of the ubiquitin-autophagosome trafficker Ref(2)p/p62 in *Gba1b^-/-^* fly brains (Figure 5E). Interestingly, Nrf2 overexpression had no significant effect on lysosomal area or ubiquitin puncta in control brains, demonstrating that the antioxidant response specifically in *Gba1b^-/-^* flies negatively impacts disease states in the brain and renal system.

**Figure 5.**
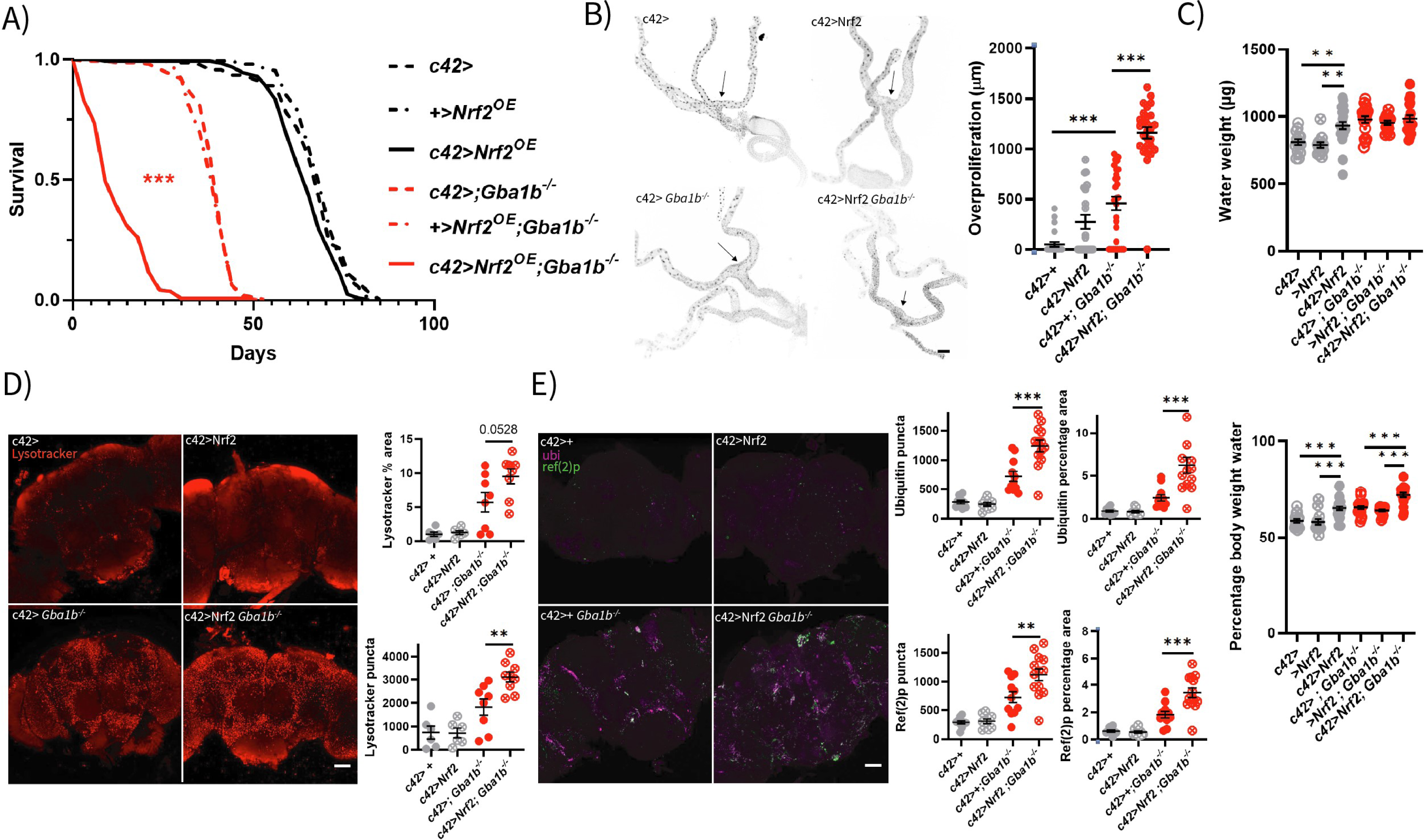
Induction of antioxidant effector Nrf2 worsens renal and neurological phenotypes in *Gba1b^-/-^*flies. A) Overexpression of Nrf2 in the MTs with the c42-Gal4 driver modestly but significantly reduces lifespan in ctrl flies and leads to a marked reduction in lifespan in *Gba1b^-/-^* flies (log-rank test: *c42>* vs *c42>Nrf2*, ***p=0.0017*; Nrf2 vs c42>Nrf2, ***p=*0.0002 ; *c42; Gba1b^-/-^ vs c42>Nrf2; Gba1b^-/-^, \*\*\**p<0.0001; *Nrf2; Gba1b^-/-^* vs *c42>Nrf2; Gba1b^-/-^*, ***p<0.0001). B) Representative 10x images and quantification of proliferating RSCs in *c42>Nrf2; Gba1b^-/-^* flies and ctrl. Overexpression of Nrf2 in the tubules leads to increased proliferation of RSCs with a significantly greater effect observed in *Gba1b^-/-^* mutants (one-way ANOVA with Tukey’s multiple comparisons test: *c42* vs *c42; Gba1b^-/-^*, ***p<0.0001; *c42>Nrf2 vs c42>Nrf2; Gba1b^-/-^, ***p<0.0001*). The arrow indicates the ureter fork. Scale bar = 100 μm. C) Water weight and percentage body water content of *c42>Nrf2; Gba1b^-/-^*flies and ctrl. Nrf2 overexpression significantly increases body water content in ctrl flies, but not in *Gba1b^-/-^* flies (one-way ANOVA with Tukey’s multiple comparisons test: *c42>* vs *c42>Nrf2*, ***p<0.0001; *Nrf2 vs c42>Nrf2, \*\*\**p<0.0001; *c42; Gba1b^-/-^* vs *c42>Nrf2; Gba1b^-/-^*, ***p<0.0001; *Nrf2; Gba1b^-/-^ vs c42>Nrf2; Gba1b^-/-^, **\*\*\****p<0.0001). D) Representative 20x images and quantification of LysoTracker^TM^ (Red)-stained brains from *c42>Nrf2; Gba1b^-/-^* flies and ctrl. Overexpression of Nrf2 in the renal tubules results in a significant increase in lysosomal puncta number in the brains of *Gba1b^-/-^* flies (one-way ANOVA with Tukey’s test: *c42; Gba1b^-/-^* vs *c42>Nrf2; Gba1b^-/-^*, **p = 0.0076). Scale bar = 50 μm. E) Representative 20x images and quantification of FK2/polyubiquitin and Ref(2)p immunostaining in the brains of *c42>Nrf2; Gba1b^-/-^* flies and ctrl. Overexpression of Nrf2 in renal tubules leads to a significant increase in polyubiquitin accumulation and FK2-positive puncta and area in *Gba1b^-/-^* fly brains (one-way ANOVA with Tukey’s test: FK2 puncta *c42; Gba1b^-/-^* vs *c42>Nrf2; Gba1b^-/-^*, ***p< 0.0001; FK2 area *c42; Gba1b^-/-^* vs *c42>Nrf2; Gba1b^-/-^*, ***p=0.0002; Ref(2)p puncta *c42; Gba1b^-/-^* vs *c42>Nrf2*; *Gba1b^-/-^* **p=0.0061, Ref(2)p area *c42*; *gba1b^-/-^* vs *c42>Nrf2*; G*ba1b^-/-^*, ***p=0.0002). Scale bar = 50 μm.

### *Gba1b^-/-^*lifespan is shortened by shifting antioxidant state at an organismal level and in renal tubules

To assess the ability of *Gba1b^-/-^* flies to respond to exogenous antioxidants, we subjected them to a diet supplemented with ascorbic acid, a vitamin C analogue previously shown to extend lifespan in healthy flies (Bahadorani et al., 2008). While ascorbic acid led to a mild, non-significant increase in lifespan in control flies, it resulted in a significant reduction in lifespan in *Gba1b^-/-^* flies, indicative of an increased sensitivity to antioxidant supplementation (Figure 6A).

**Figure 6.**
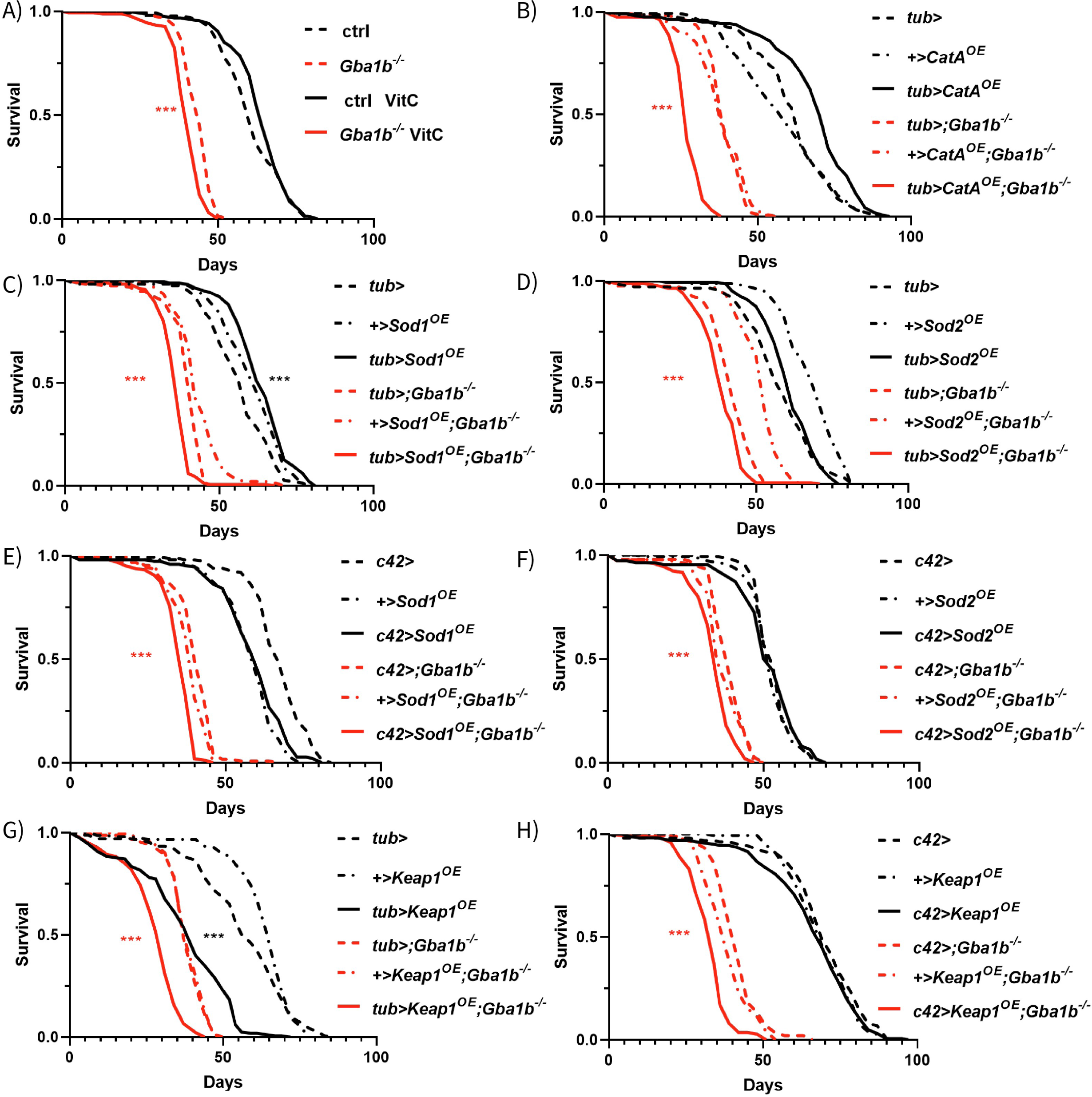
Ubiquitous or tubule-specific induction of redox dyshomeostasis shortens the lifespan of *Gba1b^-/-^*flies. A) Survival curves of ctrl and *Gba1b*^−/−^ flies raised on standard SYA food with or without 20 mM ascorbic acid (AA). AA supplementation significantly reduces lifespan in *Gba1b*^−/−^ flies (log-rank test: *Gba1b^-/-^* vs *Gba1b^-/-^* +AA ***p < 0.0001; ctrl vs ctrl *+*AA p=0.071). (B–D) Survival curves showing the effects of ubiquitous overexpression (tub-GAL4) of antioxidant enzymes *CatA* (B), *Sod1* (C), and *Sod2* (D) in *Gba1b*^−/−^ flies. All conditions resulted in significant lifespan shortening in the context of *Gba1b* loss (log-rank tests: (B) *tub>; Gba1b^-/-^* vs *tub>CatA*; *Gba1b^-/-^,* ***p<0.0001; *CatA;Gba1b^-/-^* vs *tub>CatA;Gba1b^-/-^*, ***p<0.0001. (C) *tub>; Gba1b^-/-^* vs *tub>Sod1;Gba1b^-/-^*, ***p<0.0001; *Sod1*;*Gba1b^-/-^* vs *tub>Sod1;Gba1b^-/-^*, ***p< 0.0001; (D) *tub>; Gba1b^-/-^* vs *tub>Sod2*; *Gba1b^-/-^*, ***p<0.0001; *Sod2*; *Gba1b^-/-^*vs tub>*Sod2*; *Gba1b^-/-^*, ***p<0.0001). (E–F) Lifespan analysis of *Gba1b*^−/−^ flies with *c42-GAL4* overexpression of *Sod1* (E) and *Sod2* (F), showing significant reduction in lifespan (log-rank test: *tub>*;*Gba1b^-/-^* vs *c42>Sod1*; *Gba1b^-/-^*, ***p<0.0001; *Sod1*;*Gba1b^-/-^* vs *c42>Sod1*;*Gba1b^-/-^*, ***p<0.0001. (F) *c42>;Gba1b^-/-^*vs c42>Sod2; *Gba1b^-/-^*, ***p<0.0001; *Sod2*;*Gba1b^-/-^*vs *c42>Sod2*; *Gba1b^-/-^*, ***p<0.0001). (G–H) Ubiquitous (G) and tubule (H) overexpression of the Nrf2 repressor *Keap1* in *Gba1b*^−/−^ flies significantly shortens lifespan (log-rank test: *tub>; Gba1b^-/-^* vs *tub>Keap1*;*Gba1b^-/-^*, ***p<0.0001; *Keap1*;*Gba1b^-/-^* vs *tub>Keap1*;*Gba1b^-/-^,* ***p<0.0001. (H) *c42>*;*Gba1b^-/-^* vs *c42>Keap1*; *Gba1b^-/-^*, ***p<0.0001; *Keap1*; *Gba1b^-/-^* vs *c42>Keap1*;*Gba1b^-/-^*, ***p < 0.0001).

Given that Nrf2 overexpression shortens lifespan in *Gba1b^-/-^* flies, we investigated the effects of overexpressing its downstream antioxidant targets, *Sod1*, *Sod2*, and *CatA*, both ubiquitously using the *tub-Gal4* driver and with *c42-Gal4*, which expresses in PCs. While antioxidant gene overexpression provided variable benefits in healthy control flies, ubiquitous or tubule overexpression was detrimental in *Gba1b^-/-^* flies, shortening the lifespan (Figure 6B-F), suggesting that MT and organismal responses to oxidative stress may advance disease phenotypes and contribute to systemic redox imbalance.

Furthermore, we examined the effects of PGC1α overexpression, a key regulator of mitochondrial biogenesis and an upstream activator of Nrf2. Ubiquitous overexpression of PGC1α extended lifespan in healthy controls but did not significantly impact the lifespan of *Gba1b^-/-^* flies (Figure S3C). Expression of either PGC1α or NDI, a yeast NADH dehydrogenase that bypasses mitochondrial complex I (Sanz et al., 2010), using *c42-Gal4* led to a reduction in lifespan in *Gba1b^-/-^* flies, suggesting that mitochondrial or ETC defects in tubules may not be the key driver of redox-stress in *Gba1b^-/-^*(Figure S3C, D).

Given that antioxidant activation is detrimental in *Gba1b^-/-^*flies, we investigated whether repressing the antioxidant response could have a protective effect. To this end, we expressed the Nrf2 repressor *Keap1* ubiquitously and specifically in the MTs of *Gba1b^-/-^* flies. However, *Keap1* overexpression, whether ubiquitous or using the PC driver *c42-Gal4*, significantly reduced lifespan in both *Gba1b^-/-^* flies and controls (Figure 6G, H). Taken together, our results demonstrate that flies lacking *Gba1b* experience redox dyshomeostasis and are highly susceptible to both oxidative and reductive stress.

### Parkin mutants phenocopy the renal dysfunction observed in *Gba1b^-/-^* flies

As the kidney-PD axis remains underexplored in animal models of disease, we investigated whether renal pathology is also evident in other PD models. While PD is primarily idiopathic, rare familial forms have been identified. PARK2, an E3 ubiquitin ligase, plays a crucial role in mitophagy initiation, and autosomal recessive mutations in the fly *Parkin* gene, *park*, lead to early-onset PD through mitochondrial dysfunction and subsequent redox stress (Whitworth & Pallanck, 2017). We found that loss-of-function (LOF) park mutants (*park^1/1^*) exhibited increased water weight and body water percentage (Figure 7A), which correlated with degenerative MT pathology. These mutants displayed elevated RSC proliferation (Figure 7B), increased tubule diameter, reduced mitochondrial membrane potential (Figure 7C), and a significant shift toward a more reduced mitochondrial redox state (Figure 7D), phenotypes that closely resemble those observed in *Gba1b^−/−^* fly mutants.

**Figure 7.**
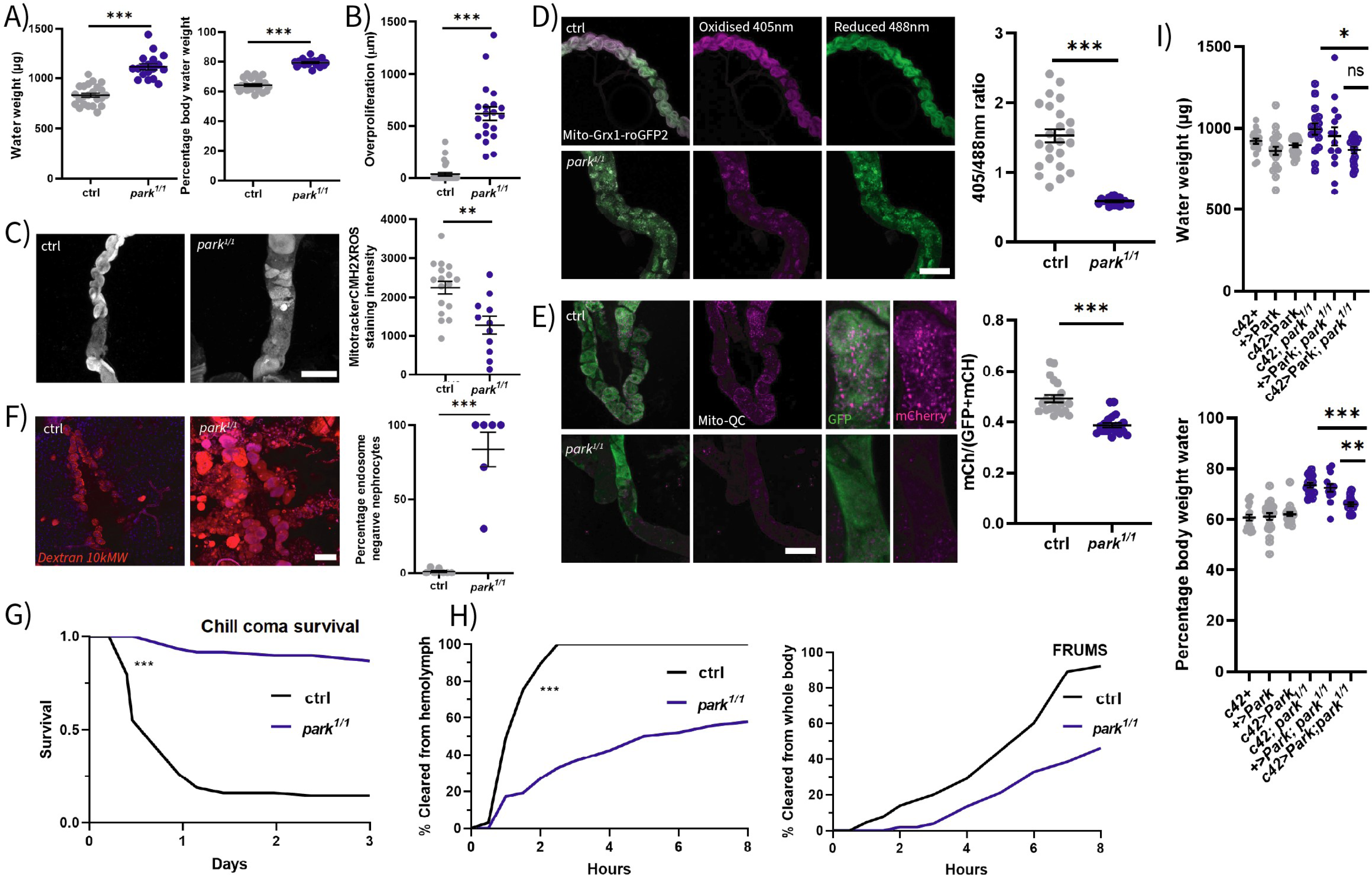
Parkin PD model flies exhibit severe renal degeneration and mitochondrial redox imbalance. (A) Water content measurements reveal significantly increased water weight and percentage body water in *park*^1/1^ flies compared to ctrl (unpaired t-test, \*\*\**p*<0.0001). (B) *park*^1/1^ flies display over proliferation of tubule RSCs in the ureter relative to ctrl (unpaired t-test, \*\*\**p*<0.0001). (C) Representative images and quantification of mitochondrial ROS (mitoROS) in the MTs show reduced mitochondrial membrane potential in *park*^1/1^ flies compared to ctrl (unpaired t-test, \*\**p*=0.0014). Scale bar = 100 µm. (D) Quantification of mitochondrial glutathione redox state using the *c42>mitoGrx1* sensor demonstrates a significant shift toward a reduced state in *park^1/1^* fly MTs (unpaired t-test, \*\*\**p*<0.0001). Scale bar = 100 µm. (E) Expression of the mitophagy reporter *c42>mitoQC* reveals a strong reduction in acidified mitolysosomes in *park^1/1^* fly tubules, indicating impaired mitophagy (unpaired t-test, \*\*\**p*<0.0001). Scale bar = 100 µm. (F) Dextran uptake assay (10×, 63×) shows a complete loss of functional nephrocytes in *park*^1/1^ flies, as quantified by the number of endocytically inactive cells (unpaired t-test, \*\*\**p*<0.0001). Scale bar = 100 µm. (G) *park*^1/1^ flies display increased survival following cold shock compared to ctrl (log-rank test, ***p=0.0007). (H) FRUMS assay reveals significantly reduced dye clearance from the hemolymph in *park*^1/1^ flies (\*\*\**p*<0.0001), though total dye clearance from the whole fly was not significantly affected (log-rank test, *p* =0.4599). (I) Overexpression of *park* in PCs (*c42*>*park*) partially rescues body water homeostasis in *park*^1/1^ flies. Water weight: *c42*; *park*^1/1^ vs *c42>park*; *park^1/1^*, \**p*=0.0471; *+>park*; *park^1/1^*vs *c42>park*; *park*^1*/1*^, *p* = 0.4201. Percentage body water: *c42; park*^1/1^ > vs *c42>park*; *park*^1*/1*^, ***p=0.0006; *+>park*; *park^1/1^*vs *c42>park*; *park*^1/1^, \*\**p*=0.0059 (one-way ANOVA with Tukey’s multiple comparisons test).

Mitophagy occurs in a spatially segregated manner in the mammalian kidney, where intense mitolysosome staining is visible in the proximal convoluted tubule, the site of high active transport demands and, consequently, elevated mitochondrial activity and turnover (Mcwilliams et al., 2016). To determine if mitophagy defects were present in *park^1/1^* mutant fly MTs, we expressed the mitophagy reporter mitoQC in PCs. This revealed a significant reduction in acidified mitolysosomes, indicating a mitophagy defect in *park^1/1^* tubules (Figure 7E).

Consistent with reduced MT function, *park1/1* nephrocytes showed a more severe phenotype than *Gba1b^-/-^*flies, with a complete loss of endocytically active nephrocytes by 28 days, as assessed by a dextran uptake assay (Figure 7F). In addition, *park^1/1^*mutants exhibited increased resistance to cold stress (Figure 7G), along with a significant reduction in renal clearance of haemolymph, as assessed by the FRUMS assay (Figure 7H).

To determine whether the effects of *park* loss were tissue-autonomous, we overexpressed *park* in *park^1/1^* mutants using the PC-driver *c42-Gal4*. This was sufficient to rescue water content, indicating that mitochondrial clearance and function are critical for maintaining tubule integrity (Figure 7I). In contrast, brain-restricted *Parkin* expression had no impact on tubule health (Figure S4A). These findings support a model in which mitophagy defects due to loss of *park* drive mitochondrial dysfunction, leading to physiological impairment and eventual tubular degeneration.

### Rapamycin alleviates disease phenotypes in the MTs of *Gba1b^-/-^* flies

To identify therapeutics with potential relevance to renal defects in our disease models, we first treated *Gba1b^-/-^* flies with rapamycin, which we previously showed extends lifespan in association with autophagy induction (Kinghorn et al 2016; Atilano et al., 2023). Rapamycin treatment significantly reduced water retention (Figure 8A), suppressed RSC over proliferation (Figure 8B) and increased FRUMS dye clearance from the body (Figure S5C, D), indicating that mTOR inhibition confers renal benefits in *Gba1b^-/-^* flies. Additionally, rapamycin normalised lipid peroxidation in *Gba1b^-/-^*MTs to control levels (Figure 8C).

**Figure 8.**
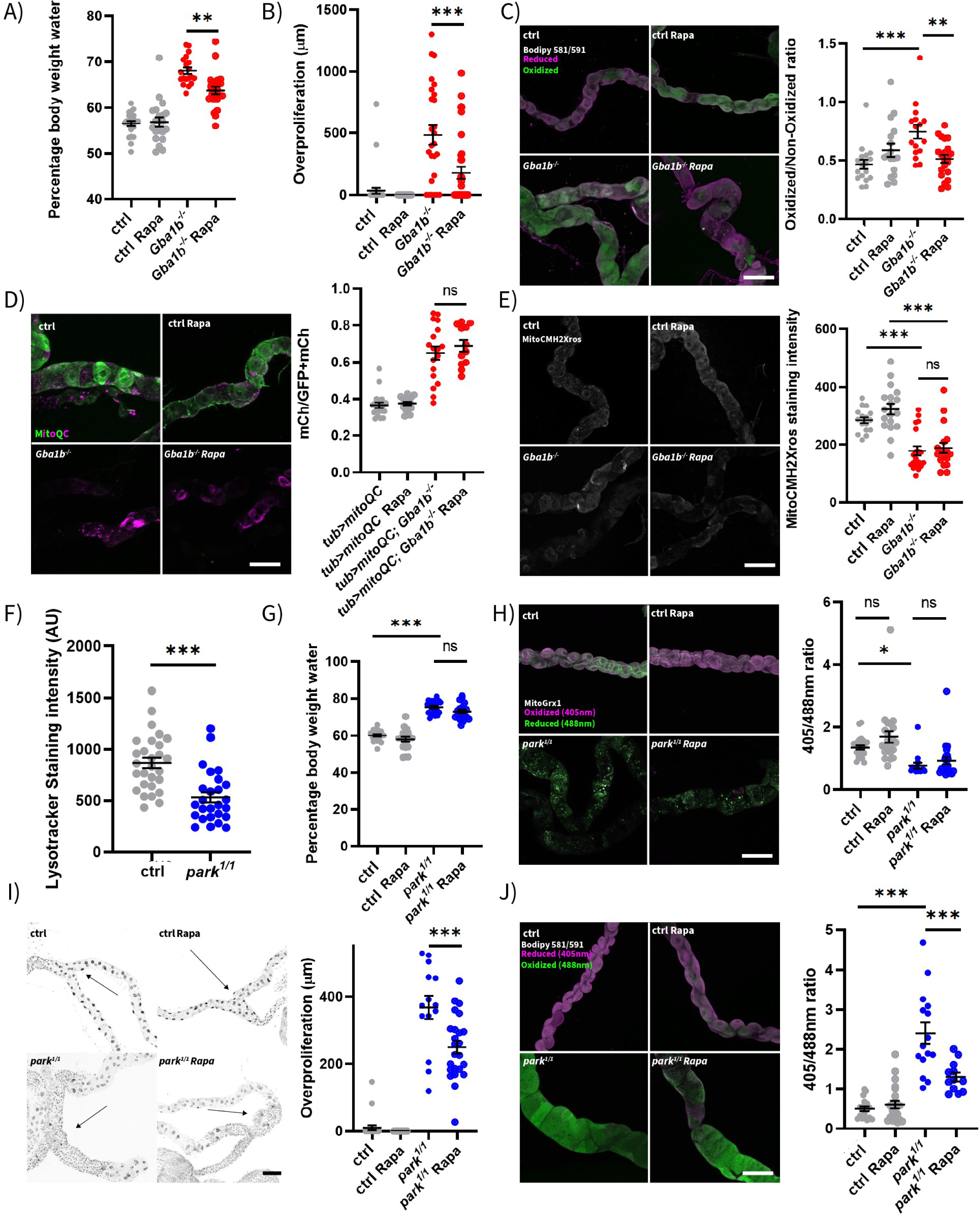
Rapamycin differentially rescues renal defects in *park^1/1^* and *Gba1b^-/-^* flies. (A) Percentage body water weight is significantly reduced in *Gba1b^-/-^*flies upon rapamycin treatment (\*\**p*=0.0017, one-way ANOVA with Tukey’s multiple comparisons test). (B) Quantification of RSC over proliferation using DAPI staining from the ureter shows a significant reduction in *Gba1b^-/-^* fly tubules following 3-week rapamycin treatment (one-way ANOVA with Tukey’s multiple comparisons test, \*\*\**p*<0.0001). (C) Lipid peroxidation, assessed using BODIPY^TM^ 581/591 (oxidised = green, reduced = magenta; excitation at 488 nm and 568 nm), is elevated in *Gba1b^-/-^* fly tubules compared to ctrl and rescued by rapamycin (one-way ANOVA with Tukey’s multiple comparisons test, \*\*\**p*<0.0001; \*\**p*=0.0042). Scale bar = 100 µm. (D) Expression of a Mitophagy reporter (tub>mitoQC) shows no significant change in mitophagy in the MTs of *Gba1b^-/-^* flies following treatment with rapamycin (one-way ANOVA with Tukey’s multiple comparisons test, *p*=0.7293). Scale bar = 100 µm. (E) Mitochondrial membrane potential visualised by MitoTracker™ Red CM-H₂XROS is significantly reduced in *Gba1b^-/-^* fly tubules compared to ctrl and is not rescued by rapamycin (one-way ANOVA with Tukey’s multiple comparisons test, \*\*\**p*<0.0001; *p*=0.9705). Scale bar = 100 µm. (F) Lysosomal content, visualised with LysoTracker™ Red, is significantly decreased in *park^1/1^* fly MTs (unpaired t-test, \*\*\**p*<0.0001). (G) Water retention is not significantly reduced by rapamycin treatment in *park^1/1^*flies (one-way ANOVA, p=0.3152). (H) Mitochondrial GSH redox state in *park^1/1^*fly MTs, visualised using c42>mitoROGFP2-Grx1, is not rescued by rapamycin (one-way ANOVA with Tukey’s multiple comparisons test, p=0.8582). Scale bar = 100 µm. (I) Rapamycin treatment reduces RSC overproliferation in *park^1/1^* fly tubules (ctrl vs *park^1/1^*, ***p<0.0001, rapa ctrl vs rapa *park^1/^*, *p<0.0109, *park^1/1^* vs rapa *park^1/1^*, ***p<0.0001, one-way ANOVA with Tukey’s multiple comparisons test). Scale bar = 100 µm. (J) BODIPY^TM^ 581/591 C11 staining shows elevated lipid peroxidation in *park^1/1^* fly tubules, which is significantly reduced by rapamycin treatment (one-way ANOVA with Tukey’s multiple comparisons test, \*\**p*<0.0001; p=0.0109). Scale bar = 100 µm.

Using tubule-specific expression of mitoQC, we found that *Gba1b^-/-^*fly MTs exhibited a reduced ratio of non-acidified to acidified mitochondria compared to control flies, suggesting that unlike *park^1/1^* tubules, mitophagy remains functional and potentially overactive. Moreover, the mitoQC ratio in *Gba1b^-/-^* fly MTs was not affected by rapamycin treatment (Figure 8D). The decreased mitochondrial membrane potential of *Gba1b^-/-^* MTs was not rescued by rapamycin treatment (Figure 8E). This indicates that some aspects of mitochondrial dysfunction in *Gba1b^-/-^* flies are independent of mTOR-regulated pathways.

In contrast to the MTs of *Gba1b^-/-^* flies, *park^1/1^*fly tubules displayed decreased lysosomal content (Figure 8F), consistent with a block in mitophagy. Rapamycin failed to rescue the elevated water weight (Figure 8G), or mitochondrial redox state (Figure 8H) in *park^1/1^* flies, indicating that renal dysfunction in this model may stem from mitophagy defects that are unresponsive to mTOR inhibition. However, rapamycin did rescue both ureter-associated RSC over proliferation and lipid peroxidation in *park^1/1^* fly tubules (Figures 8I, J). Together, these findings suggest that although both *Gba1b^-/-^* and *park^1/1^* mutant flies exhibit pronounced renal defects, the underlying mechanisms differ. *park^1/1^* fly mutants display a mitophagy block and renal dysfunction that cannot be reversed by rapamycin, whereas the renal pathology in *Gba1b^-/-^* flies is influenced by rapamycin, potentially through altered systemic lipid metabolism.

## Discussion

Despite growing epidemiological evidence linking renal disease to neurodegenerative disorders (Meléndez-Flores & Estrada-Bellman, 2021), the mechanistic and directional relationships between these conditions remain unresolved. Here, we identify progressive renal dysfunction as a previously underappreciated feature in *Drosophila* models of GD and PD. Flies lacking either the main *GBA1* orthologue, *Gba1b,* or the mitophagy regulator *Parkin, park,* exhibit age-dependent degeneration of the renal system. This is characterised by disorganised MTs, impaired nephrocyte function, redox imbalance, and lipid accumulation. These renal impairments contribute to organismal decline, including fluid imbalance, ionic hypersensitivity, and worsening neurodegeneration.

A key finding is that *Gba1b*-deficient flies exhibit a collapse of redox homeostasis that does not align with classical oxidative stress. Redox-sensitive probes revealed lipid peroxidation and cytosolic oxidation alongside modest mitochondrial reduction in *Gba1b*⁻^/^⁻ MTs. This divergence was further supported by opposite responses to redox-targeted interventions. In *Gba1b*^⁻/^⁻ flies, overexpression of Nrf2 or antioxidant enzymes (SOD1, SOD2, CatA), and supplementation with ascorbic acid, all led to worsened survival. These results suggest a critical loss of redox buffering capacity, where both oxidative and reductive shifts are deleterious. This redox fragility sets *Gba1b*-related pathology apart from conventional oxidative stress models (Dias et al., 2014) and echoes recent findings that both oxidative and reductive stress can impair renal function in *Drosophila* (Burbridge et al., 2021). These observations underscore the potential risks of unselective antioxidant therapies in *GBA1*-related diseases and suggest that redox tone may serve as a biomarker of tissue vulnerability and therapeutic responsiveness.

In contrast to *Gba1b^-/-^* flies, those lacking *park* benefit from antioxidant pathway activation (Whitworth et al., 2005), consistent with oxidative stress as a key driver of pathology. This model-specific disparity underscores the need for tailored therapeutic approaches. Clinically, mitophagy has been shown to protect against renal injury and nephrotoxicity (Tang et al., 2018; Wang et al., 2018), reinforcing *parkin*’s role in renal health.

Notably, the source of renal damage differs between models. In *park*^1/1^ flies, renal defects are tissue-autonomous: tubule-specific re-expression of *Parkin* restores water regulation. Conversely, *Gba1b* is not expressed in the renal system, pointing to a non-cell-autonomous mechanism of dysfunction. This suggests that systemic factors, such as circulating glycosphingolipids or a loss of extracellular vesicle-mediated delivery of GCase, may mediate renal toxicity (Thomas et al., 2018). Supporting this hypothesis, sphingolipid accumulation is a hallmark of GD, and lipid dysregulation has been directly implicated in renal pathology. For example, mutations in the ceramide synthase gene *CERS2* cause renal dysfunction in mice (Imgrund et al., 2009). Moreover, ceramide buildup induces podocyte apoptosis (Yoo et al., 2015) and *Drosophila sply* mutants, deficient in sphingosine-1-phosphate lyase, also display nephrotic-like phenotypes with slit diaphragm defects (Lovric et al., 2017). It should be noted, however, that these are developmental and static in nature, unlike the progressive degeneration in *Gba1b*⁻^/^⁻ flies.

Mitochondrial analysis further revealed that *Gba1b* deficiency impairs mitochondrial morphology, reduces membrane potential, and limits NADH availability, all hallmarks of dysfunction seen in *Gba1*-deficient mammalian neurons (Osellame et al., 2013). However, mitophagy appears preserved in *Gba1b*⁻^/^⁻ MTs, in contrast to *park*^1/1^ flies. A reduction in intact mitochondria in both models suggests that excessive degradation may contribute to renal stress, albeit through divergent upstream mechanisms.

Strikingly, rapamycin, a known mTOR inhibitor and autophagy inducer (Bjedov et al., 2010), rescued renal structure and function in *Gba1b*^⁻/^⁻ flies, including water retention, RSC proliferation, and lipid peroxidation. This aligns with findings that mTOR overactivation contributes to podocyte damage in glomerular disease (Godel et al., 2011). These effects occurr despite unaltered mitophagy, pointing toward alternative mechanisms such as improved lipid handling or redox buffering. In contrast, rapamycin had limited effects on renal function in *park¹/¹* flies, highlighting mechanistic divergence between lysosomal and mitochondrial models. In GD, where lysosomal clearance is impaired, rapamycin may enhance autophagic flux to bypass the enzymatic block (Atilano et al., 2023).

Together, these findings establish renal degeneration as a driver of systemic decline in *Drosophila* models of GD and PD. Recent evidence for a brain-kidney axis in PD, where α-synuclein accumulates in renal tissue and may retrogradely propagate to the brain under renal failure (Yuan et al., 2025), supports the relevance of the renal system to PD. We propose that renal oxidative stress could influence such inter-organ signalling, with potential feedback on neurodegenerative cascades. Future validation of these mechanisms in mammalian models will be essential to define translational relevance.

If confirmed, our findings could inform new biomarker strategies and therapeutic targets for *GBA1* mutation carriers and other at-risk groups. Maintaining renal health may represent a modifiable axis of intervention in neurodegenerative disease. Our results also reinforce the need for precision in redox modulation, particularly where antioxidant therapies may cause harm. Finally, the opposing effects of rapamycin in *Gba1b* and *park* models demonstrate that effective treatments must account for the underlying molecular defect. Renal imaging or biomarker assays in *GBA1* mutation carriers may provide a means to monitor disease progression or therapeutic response. Identifying redox fragility and mTOR sensitivity as actionable parameters opens new avenues for screening in both invertebrate and vertebrate systems.

In conclusion, we uncover a critical and previously overlooked role for the renal system in GD and PD pathogenesis. We demonstrate that redox dyshomeostasis, renal degeneration, and mTOR-responsive pathways are central features of disease in *Gba1b*-deficient flies, and mechanistically distinct from those in *Park* mutant models. These findings support a broader, integrated view of neurodegenerative disease and pave the way for organ-specific interventions in disorders long considered purely neurocentric. Renal health, redox homeostasis and autophagy status emerge as key modulators of disease trajectory, offering a new conceptual and therapeutic framework beyond the brain.

## Acknowledgements

We thank all members of the Kinghorn lab and the UCL Institute of Healthy Ageing, Michael Duchen and Antonella Spinazzola for critical comments and Shu Yi Koay for early contributions to the project. This work was supported by the Wellcome Trust (Wellcome Career Development Awards to K.J.K., 214589/Z/18/Z and 309250/Z/24/Z), funding from the Rosetrees Trust (K.J.K. and A.H. M701 and M701A), the MRC (K.J.K and A.H. MR/X011070/1), and the Vivensa Foundation (K.J.K. PF2302/8).

## SUPPLEMENTAL FIGURE LEGENDS

**Supplemental Figure 1**

S1A) Quantification of NAD+/NADH ratios in whole ctrl and *Gba1b^-/-^*flies (unpaired t-test NAD+ ctrl vs *Gba1b^-/-^*, p=0.7770; NADH ctrl vs *Gba1b^-/-^*, p=0.0208*). S1B) Example image showing the localisation of *Gba1b* expression within *Drosophila* MTs. *Gba1b^CRIMIC^>mCherry.nls*, DAPI (Blue) and mCherry (Red). Scale-bar = 100µm.

**Supplemental Figure 2.**

S2A) Drinking assay using differential fly weights from a starved mean, demonstrating a significant decrease in water ingestion by *Gba1b^-/-^* flies compared to ctrl (unpaired t-test, *p=0.0057). S2B) Drinking assay by weight differential of a water source, demonstrating no significant difference in water ingestion in *Gba1b^-/-^* flies compared to ctrl (unpaired t-test, p=0.50). S2C) Ctrl and *Gba1b^-/-^* fly water weight across a desiccation time course (two-way-ANOVA, ctrl vs *Gba1b^-/-^*\*\*\*p<0.0001, 0hr vs 5hr vs 10hr ***p<0.0001). S2D) Ctrl and *Gba1b^-/-^* fly water weight across a natriuresis assay (one-way ANOVA with Tukey’s multiple comparison’s test. Ctrl SYA vs ctrl NaCl, ***p<0.0001; ctrl SYA vs ctrl NaCl + H_2_O, p=0.9748; ctrl NaCl vs ctrl NaCl + H_2_O, ***p<0.0001; *Gba1b^-/-^* SYA vs *Gba1b^-/-^* NaCl, ***p<0.0001; *Gba1b^-/-^* SYA vs *Gba1b^-/-^* NaCl + H_2_O, p=0.1299; *Gba1b^-/-^* NaCl vs *Gba1b^-/-^* NaCl + H_2_O, **p=0.0041) S2E) Ctrl and *Gba1b^-/-^* fly water weight before and after chill coma stress assay (two-way ANOVA ctrl vs *Gba1b^-/-^,* ***p<0.0001, 0 hr vs 10 hr, p=0.8536)

**Supplemental Figure 3**

S3A) Example images (20x) and quantification of peroxisomes of ctrl and *Gba1b^-/-^* MTs. expressing tub>YFP:PTS. Quantification of mean peroxisome area (µm^2^) (unpaired t-test, p=0.0322), percentage of MT that is peroxisome +ve (unpaired t-test, **p=0.0019) and number of peroxisomal per µm^2^ (unpaired t-test, p***=0.0003), All quantification was conducted on segmented max projection z-stacks. S3B) Example images (20x) of immunostaining for Sod1 (red; unpaired t-test, **p=0.0014) and CatA (red) (unpaired t-test, p=0.0801) in control and *Gba1b^-/-^* MTs. Scale-bar = 50µm. S3C) Survival curve for *tub>PGC1α; Gba1b^-/-^* (log-rank test, *tub>* vs *tub> PGC1α,* ***p<0.0001; *> PGC1α*; vs *tub> PGC1α,* ***p<0.0001, *tub>; Gba1b^-/-^* vs *tub> PGC1α; Gba1b^-/-^* ***p<0.0001; *> PGC1α; Gba1b^-/-^* vs *tub> PGC1α; Gba1b^-/-^*, ***p<0.0001). S3D) Survival curve for *c42> PGC1α; Gba1b^-/-^* (Log-rank test, *c42>; Gba1b^-/-^* vs *c42> PGC1α; Gba1b^-/-^*, ***p<0.0001; *> PGC1α; Gba1b^-/-^* vs *c42> PGC1α; Gba1b^-/-^*, ***p<0.0001). S3E) Survival curve for *c42>Ndi; Gba1b^-/-^* (log-rank test, *c42>; Gba1b^-/-^* vs *c42>Ndi; Gba1b^-/-^*, ***p<0.0001; *>Ndi; Gba1b^-/-^* vs *c42>Ndi; Gba1b^-/-^*, ***p<0.0001).

**Supplemental Figure 4**

S4A) Example images and quantification of DAPI-stained nuclei proliferation from the ureter fork for elav>*park*; *park^1/1^*and ctrl. (Elav>+; *park^1/1^* vs elav>park; *park^1/1^*, p=0.0743, one-way ANOVA with Tukey’s multiple comparisons test). Scale-bar = 100µm. S4B) Climbing assay time course for *klf15^-/-^; park^1/1^*demonstrating no additive effect of nephrocyte ablation on climbing phenotypes (age-effect, **p=0.0002; genotype effect, ***p<0.0001; ctrl vs *klf15^-/-^*p=0.1735; ctrl vs *park^1/1^*, ***p<0.0001, *park^1/1^* vs *klf15^-/-^ park^1/1^*, ***p=0.0003, two-way ANOVA with Tukey’s multiple comparisons test).

**Supplemental Figure 5**

S5A) Example images corresponding to figure 8B), ctrl and *Gba1b^-/-^* 3-week-old tubules treated with rapamycin and stained with DAPI. Scale-bar = 100µm. Arrow denotes the ureter fork. S5B) Example images corresponding to figure 8F), ctrl and *park^1/1^* 3-week-old tubules stained with LysoTracker^TM^, Scale-bar = 100µm. S5C) Quantification of FRUMS assay clearance from the hemolymph in ctrl and *Gba1b^-/-^* (ctrl vs *Gba1b^-/-^*, ***p<0.0001, log-rank test; ctrl vs ctrl rapa, p=0.666, log-rank test; *Gba1b^-/-^*vs *Gba1b^-/-^* rapa, p=0.0005, log-rank test). S5D) Quantification of FRUMS assay clearance from the whole body in ctrl and *Gba1b^-/-^* flies (ctrl vs *Gba1b^-/-^* cleared from haemolymph, ***p<0.0001, log-rank test; ctrl vs ctrl rapa p=0.561, log-rank test; *Gba1b^-/-^*vs *Gba1b^-/-^* rapa, **p=0.0038, log-rank test).

