## Supplementary figures and images for "Redox Dyshomeostasis Links Renal and Neuronal Dysfunction in *Drosophila* Models of Gaucher and Parkinson’s Disease"

### Figure S1

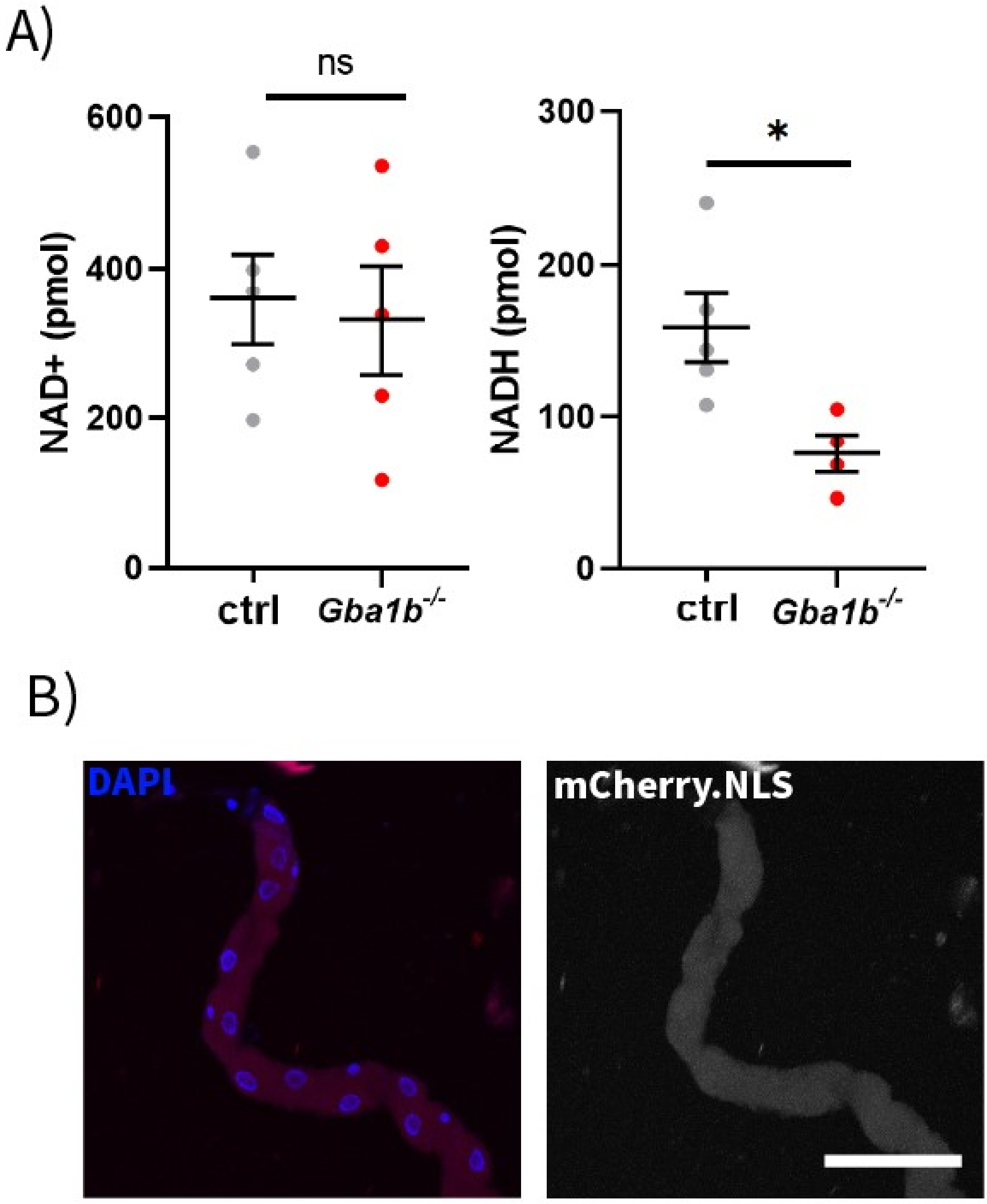

### Figure S2

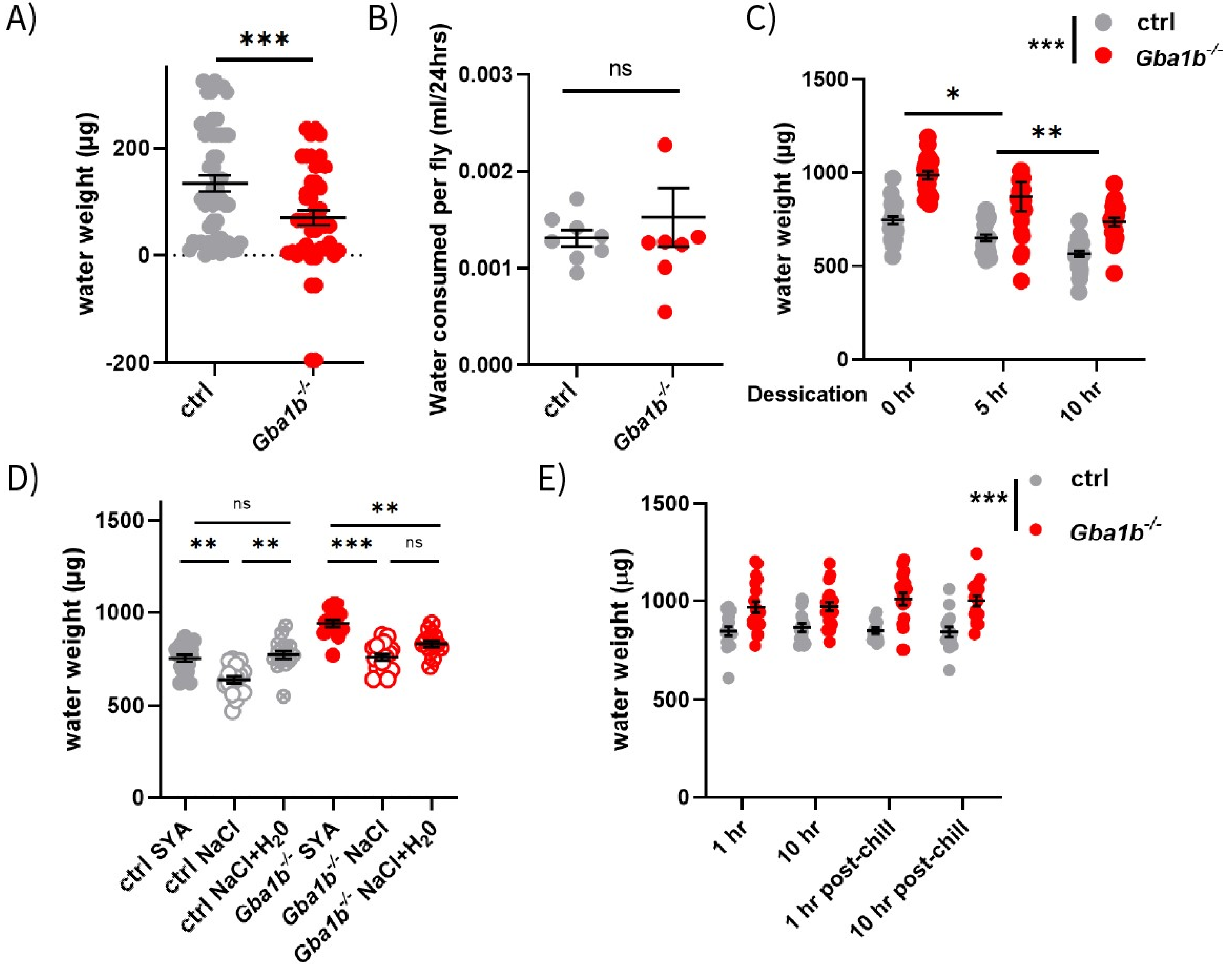

### Figure S3

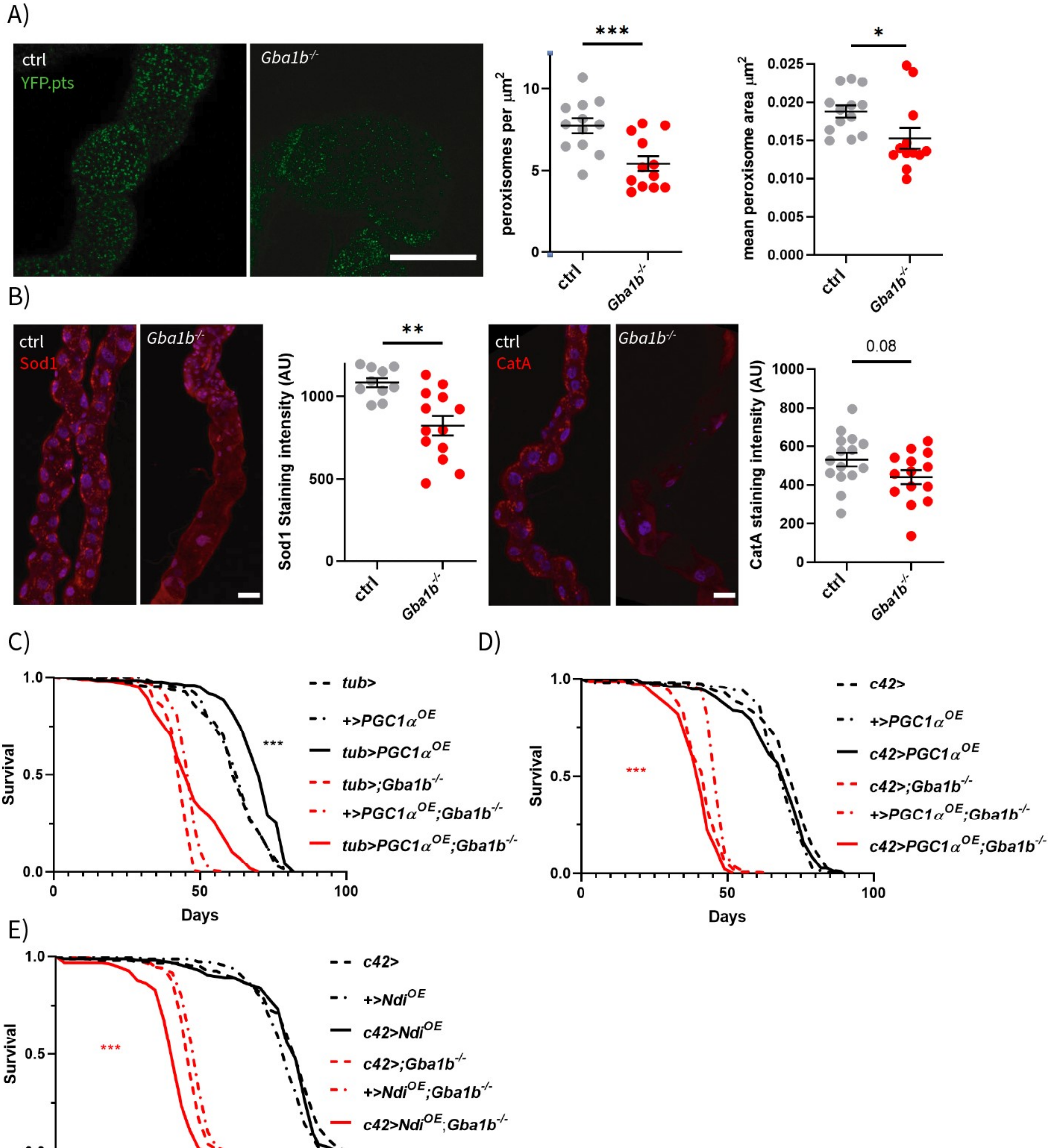

### Figure S4

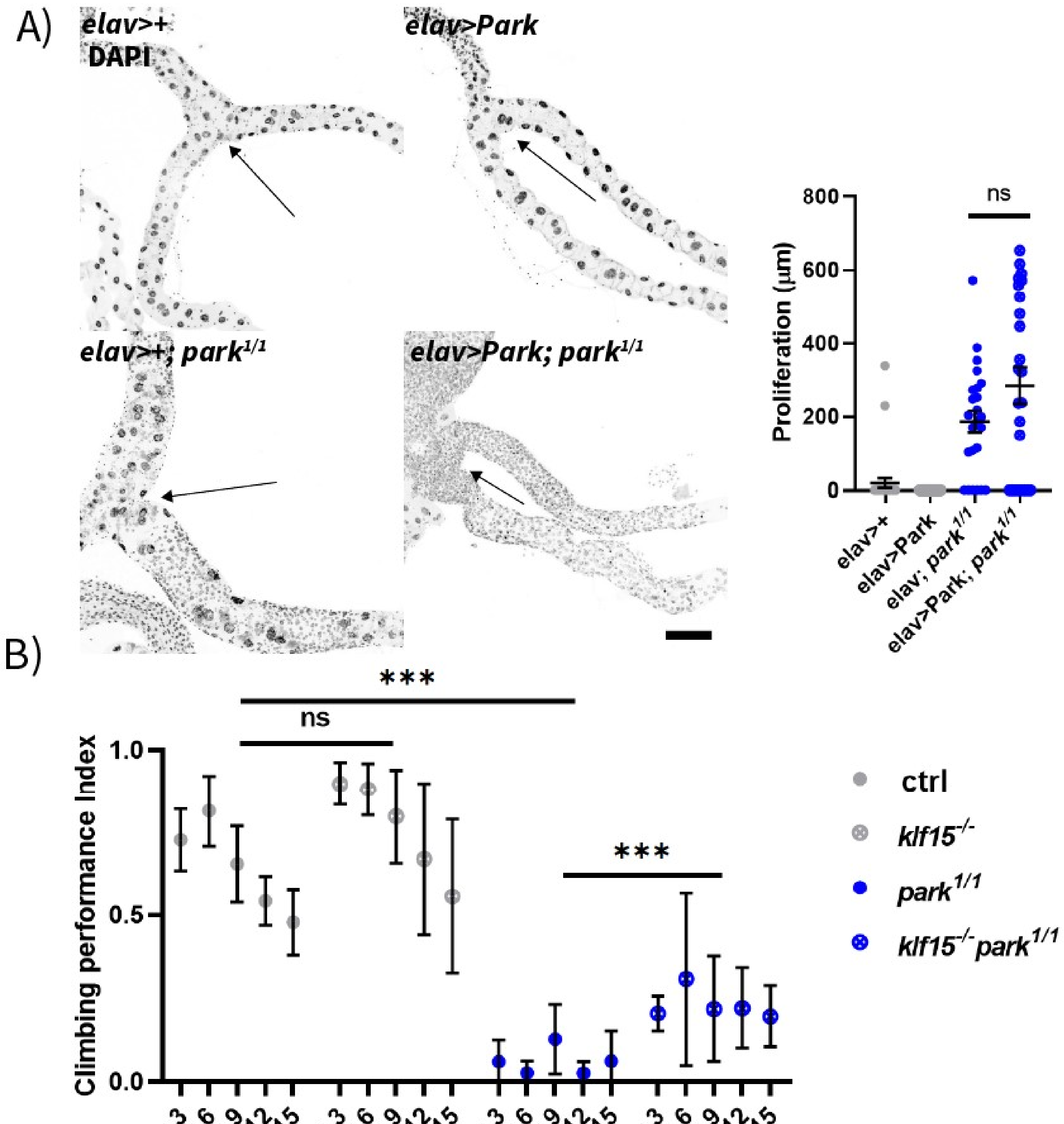

### Figure S5

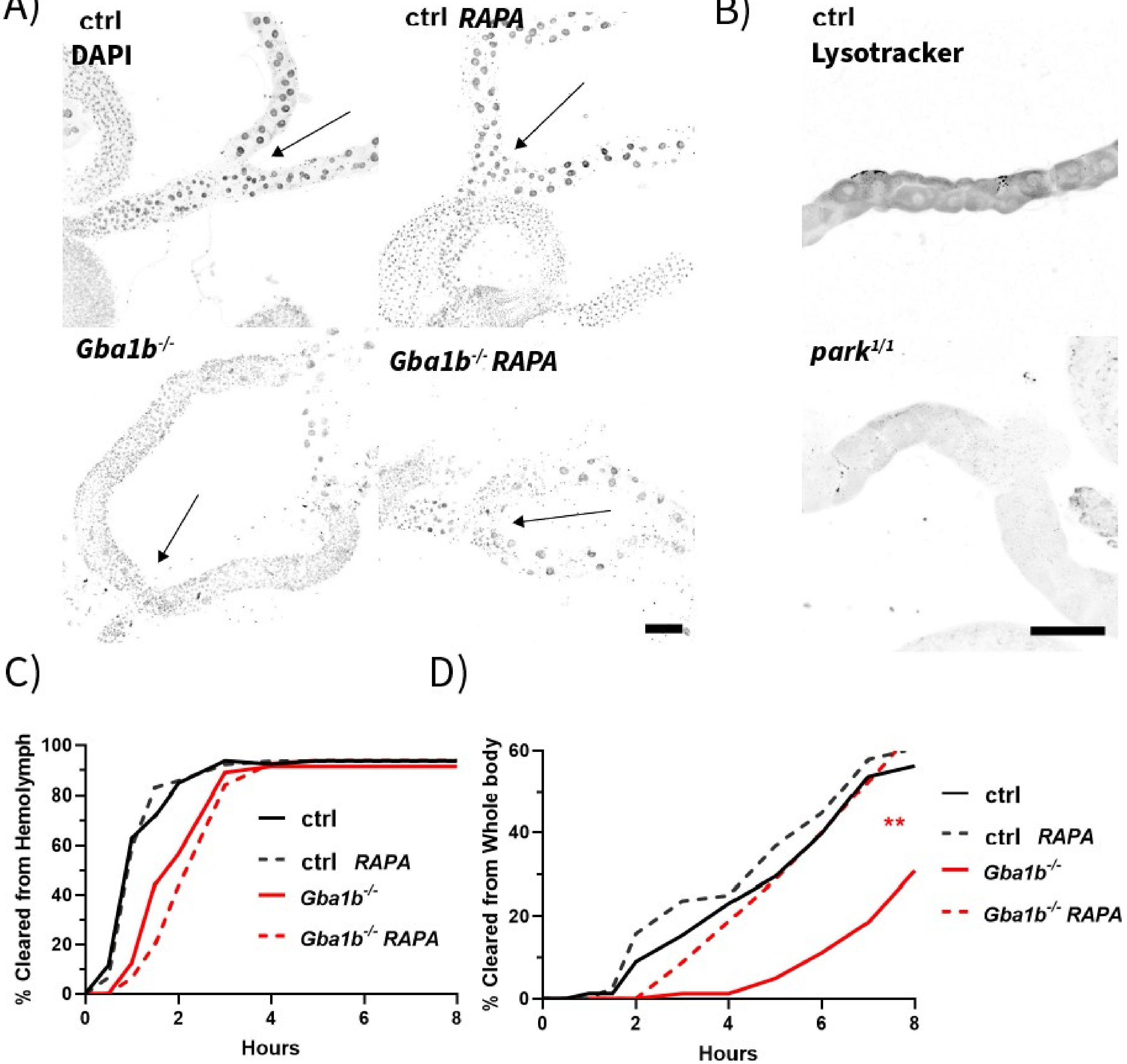
